# Multiple *FGFR1* mutations modulate tumorigenic mechanisms in glioneuronal tumors

**DOI:** 10.1101/2025.05.27.654799

**Authors:** Jacopo Boni, Míriam Fernández-González, HyeRim Han, Carla Roca, Cassandra J. Wong, Cristina Rioja, Clara Nogué, Leticia Manen-Freixa, Jonathan Boulais, Endika Torres-Urtizberea, Antonio Gomez, Martin Hasselblatt, Roger Estrada-Tejedor, Albert A. Antolin, Islam E. Elkholi, Nada Jabado, Jean-François Côté, Anne-Claude Gingras, Barbara Rivera

**Affiliations:** Bellvitge Biomedical Research Institute (IDIBELL), Avinguda de la Granvia de l’Hospitalet 199, 08908 L’hospitalet de Llobregat, Barcelona, Spain; Universitat de Barcelona (UB), Gran Via de les Corts Catalanes 585, 08007 Barcelona, Spain; Lunenfeld-Tanenbaum Research Institute, Mount Sinai Hospital, Sinai Health System, Toronto, Ontario, Canada; Institut de Recherches Cliniques de Montréal (IRCM), Montreal, Quebec, H2W 1R7, Canada; Grup de Química Farmacèutica, IQS School of Engineering, Universitat Ramon Llull, Via Augusta 390, E-08017 Barcelona, Spain; Center for Cancer Drug Discovery, The Division of Cancer Therapeutics, The Institute of Cancer Research, London, UK; Program Against Cancer Therapeutic Resistance (ProCURE), Catalan Institute of Oncology (ICO)/IDIBELL, 08908, L’Hospitalet de Llobregat, Barcelona, Spain; Department of Biosciences, Faculty of Sciences and Technology (FCT), University of Vic - Central University of Catalonia (UVic-UCC), Vic, 08500, Barcelona, Catalonia, Spain; Institute of Neuropathology, University Hospital Münster, 48149 Münster, Germany; Department of Human Genetics, McGill University, Montreal, QC, Canada; Department of Medicine, Université de Montreal, Montreal, QC, Canada; Department of Anatomy and Cell Biology, McGill University, Montreal, QC, Canada; Department of Molecular Genetics, University of Toronto, Toronto, ON, Canada; Lady Davis Institute for Medical Research, Segal Cancer Centre, Jewish General Hospital, 3755 Chemin de la Côte Sainte-Catherine, Montreal, Quebec, H3T 1E2, Canada; Gerald Bronfman Department of Oncology, McGill University, Montreal, Quebec, H4A 3T2, Canada

**Keywords:** FGFR1, multiple mutations, glioneuronal tumors, cell models, modulatory mechanisms

## Abstract

*FGFR1* genetic alterations are associated with human diseases, including brain tumors. We reported multiple *FGFR1* mutations in familial and sporadic cases of low-grade glioneuronal tumors, suggesting intrinsic mechanisms of selective pressure toward *FGFR1* multiple events arising in the context of a quiet genome. To decipher the molecular mechanisms triggered by multiple *FGFR1* mutations, we have mapped the proximal interactome of wild-type, single-and double-mutant FGFR1 proteins through a BioID-MS approach. Our data reveals novel oncogenic functionality for the two hotspots N546K and K656E, linked to evasion of lysosomal degradation. We identified a modulatory role played by the susceptibility variant R661P, which hampers the oncogenic potential of both hotspot mutations by rescuing receptor degradation and reducing N546K affinity for the downstream effector PLCγ. The R661P variant alone abolished the self-renewal capacity of oligodendroglioma cells and showed downregulation of genes involved in neurodevelopment and neuro-glial cell fate decisions, both aspects overcome in the double mutants. This study sheds light on the oncogenic effects associated with *FGFR1* alterations and their recurrence in low-mutation burden and therapy naïve tumors.

## Introduction

Fibroblast growth factor receptors (FGFRs) are a highly conserved family of receptor tyrosine kinases (RTKs) playing fundamental functions during organ development and tissue homeostasis (Xie *et al*, 2020). Hence, genetic alterations in FGFR genes have been associated with a broad spectrum of human diseases. Constitutional pathogenic variants in *FGFR1* have been linked to the etiology of several developmental disorders, which comprise Pfeiffer syndrome, osteoglophonic dysplasia, Kallmann syndrome, and Hartsfield syndrome, among others (Dhamija & Babovic-Vuksanovic, 1993; Jarzabek *et al*, 2012; Vogels & Fryns, 2006; White *et al*, 2005). On the other hand, somatic alterations such as *FGFR1* gene amplification, duplication of the tyrosine kinase domain, gene fusions and hotspot single-nucleotide variants (SNV) have been found enriched in many types of solid tumors, including lung squamous cell carcinoma, urothelial carcinoma and glioma (Helsten *et al*, 2016).

Our group and others have described *FGFR1* genetic variants linked to the etiology of hereditary and sporadic forms of early-onset, low-grade glioneuronal tumors (LGGNTs). Specifically, duplication of the kinase domain, gene fusions and point mutations were identified as driver events in pilocytic astrocytoma, rosette-forming glioneuronal tumors (RGNTs) and other LGGNT subtypes (Engelhardt *et al*, 2022; Jones *et al*, 2013; Qaddoumi *et al*, 2016; Rivera *et al*, 2016; Sievers *et al*, 2019; Zhang *et al*, 2013). In particular, a high proportion of pediatric epileptogenic LGGNTs, defined as dysembryioplastic neuroepithelial tumors (DNETs), harbor oncogenic *FGFR1* alterations (Qaddoumi *et al*., 2016; Rivera *et al*., 2016). Histologically, these tumors are characterized by both neuronal and oligodendroglia-like elements (Daumas-Duport, 1993; Louis *et al*, 2021). Our study identified a hereditary form of DNETs with a novel *FGFR1* missense variant (R661P) mapping in the tyrosine kinase domain. Notably, additional hotspot *FGFR1* somatic mutations (either N546K or K656E) were detected *in cis* to the R661P variant in the tumors of carrier individuals (Rivera *et al*., 2016). Although double mutations in driver genes are rare events, the pattern of multiple, co-occurring single-point mutations *in cis* (hereafter referred as “multiple”) in *FGFR1* gene was validated in sporadic cases. These findings point to mechanisms of selective pressure that promote the accumulation and positive selection of multiple mutational events in *FGFR1* during glioneuronal tumor formation. This occurrence of intragenic multiple mutations in low-grade, low mutation-burden and predominantly therapy-naïve tumors, represents a striking and unexplained phenomenon.

*FGFR1* N546K and K656E oncogenic variants have been reported in postzygotic mosaicism as the genetic cause of a rare neurocutaneous condition known as Encephalocraniocutaneous lipomatosis (ECCL) (Bennett *et al*, 2016); however neither variant has so far been reported in the germline, suggesting embryonic lethality. ECCL has been classified as one of the RASopathies, a group of developmental disorders characterized by germline or mosaic mutations in genes that lead to constitutive activation of the RAS/MAPK pathway. Additional *FGFR1* missense mutations *in cis* have been reported in low-grade gliomas (LGGs) developed by ECCL individuals classified as midline pilocytic astrocytomas (Bennett *et al*., 2016; Valera *et al*, 2018), providing additional evidence for a driver role of multiple *FGFR1* mutations in LGGs/LGGNTs.

Molecularly, the two hotspot mutations have been linked to increased basal activation and ectopic expression has been shown to induce transformation and hyperproliferative phenotypes in mammalian cell lines (Cimmino *et al*, 2022; Hart *et al*, 2000; Lew *et al*, 2009; Yoon *et al*, 2004), indicating an acquired oncogenic potential by the mutated receptor. Despite this data, the precise pro-tumorigenic molecular mechanisms in glial cells remain elusive. Moreover, to date no research work has investigated how this tumorigenic activity is affected by additional *FGFR1* mutations *in cis* and why they are positively selected in LGGNTs (mostly low-mutation burden and therapy-naïve tumors). In the present study, we employ complementary models and experimental strategies, including a Proximity-dependent Biotin Identification (BioID)-based proximal interactome profiling and CRISPR-engineered glioma cell lines, to investigate how secondary hits in the oncogene *FGFR1* modulate cellular effects and oncogenic drive mediated by hotspot mutations.

Overall, our findings provide mechanistic evidence on the driver effects exerted by multiple mutations in *FGFR1* gene and shed new light on tumorigenic mechanisms during early-onset brain tumor formation.

## Results

### Hotspots and multiple *FGFR1* mutations are recurrent in brain tumors

We queried the GENIE database (de Bruijn *et al*, 2023) (https://genie.cbioportal.org/) to define the spectrum of tumors harboring at least one of the two hotspot variants N546K and K656E. These two amino acid changes represent the most frequent event among all the missense variants identified for the residues Ans546 (213/244) and Lys656 (121/137) (Appendix, Table S1). High specificity for brain tumors was confirmed for both variants (Fig. 1A, Table 1; χ^2^ test, p < 0.001). Among these cases (N546K or K656E mutated), we reviewed tumors with multiple hits in *FGFR1* and found that they were exclusive to brain tumor types, with the hotspot K656E being the one more frequently accompanied by additional *FGFR1* hits (Fig. 1A, Table 2). Secondary *FGFR1* variants (“secondary” hereafter referred as any *FGFR1* missense variant other than N546K or K656E identified in samples with multiple *FGFR1* variants) have been specified in Appendix Fig. S1. By examining tumor grade of hotspot-positive brain tumor types, we found that multiple *FGFR1* alterations are found in similar rates in both low-grade (grades 1-2) and high-grade (grades 3-4) cases, with a higher number of double-mutant cases identified in low-grade types (Fig. 1B, Table 3). Regarding the timing of occurrence and clonal selection of *FGFR1* multiple mutational events, allele frequencies do not provide enough information on whether “secondary” variants appear before or after the occurrence of the oncogenic hit. Nevertheless, we can infer hypothesis on the timing from patients with genetic disorders caused by germline and mosaic forms of *FGFR1* variants, which indicate that both scenarios (“secondary” hits appearing first as in familial DNETs, or hotspots appearing first as in ECCL) are possible (Fig. 1C).

**Figure 1.**
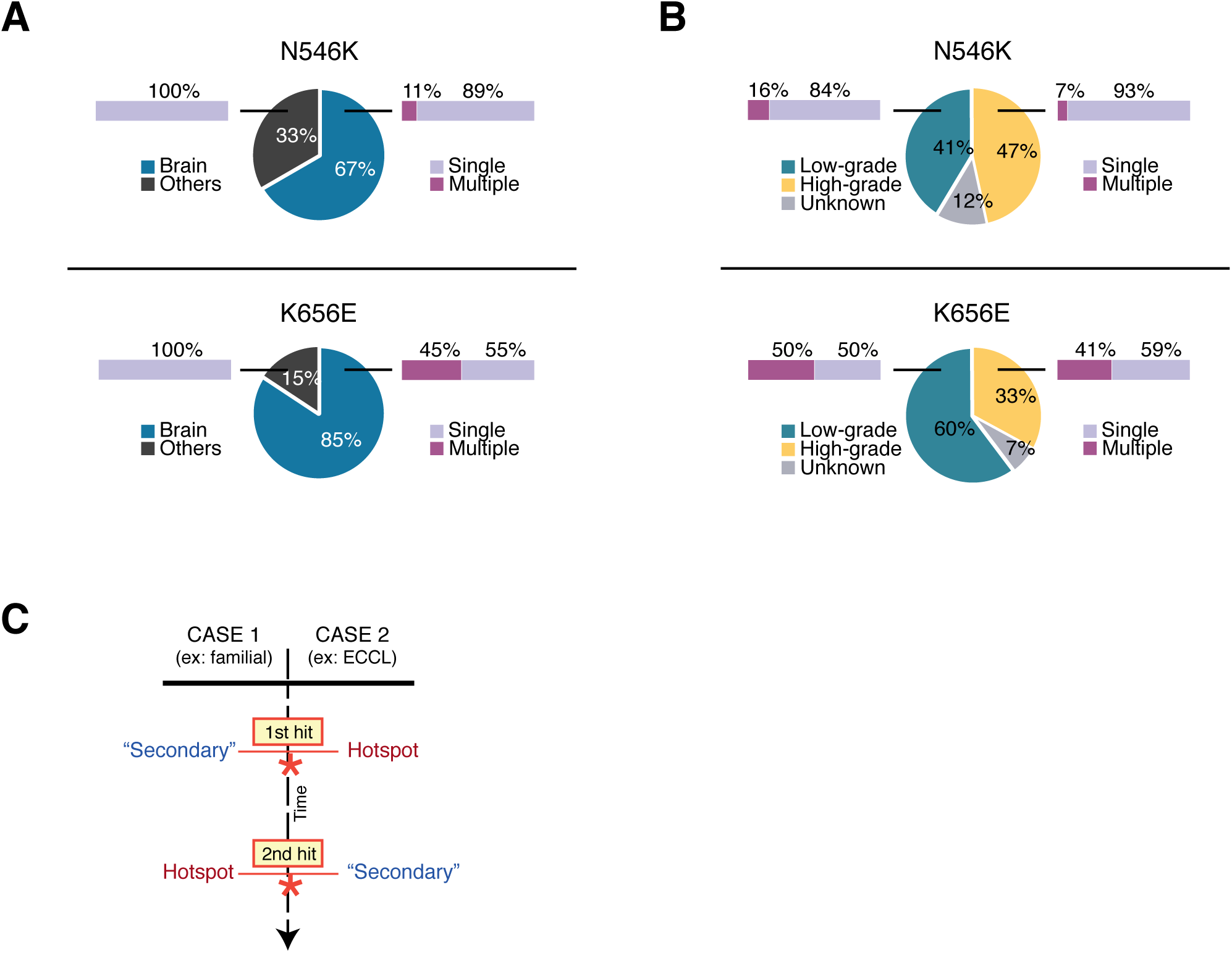
Genotype-phenotype analysis of *FGFR1* hotspots N546K and K656E, alone or with secondary hits. **A)** Distribution of N546K (n = 213) and K656E (n = 121) tumors comparing Brain vs no-brain, or “Others” tumor types. Numbers are specified in Table 1. Percentage of cases presenting only the hotspot (single) vs hotspot + other hits in *FGFR1* (multiple) are indicated for each group. Both categories (single and multiple) are significantly associated with brain tumor types (Chi-squared test, p value < 0.0001). **B)** Distribution of N546K and K656E brain tumors comparing low-grade (grade 1-2) vs high-grade (grade 3-4) tumor types. Percentage of cases presenting only the hotspot (single) vs hotspot + other hits in *FGFR1* (multiple) are indicated for each group. **C)** Timeline of acquisition of hotspot (either N546K or K656E) and “secondary” hits. The two possible scenarios with respective examples (familial case and ECCL patients) are indicated.

**Table 1.**
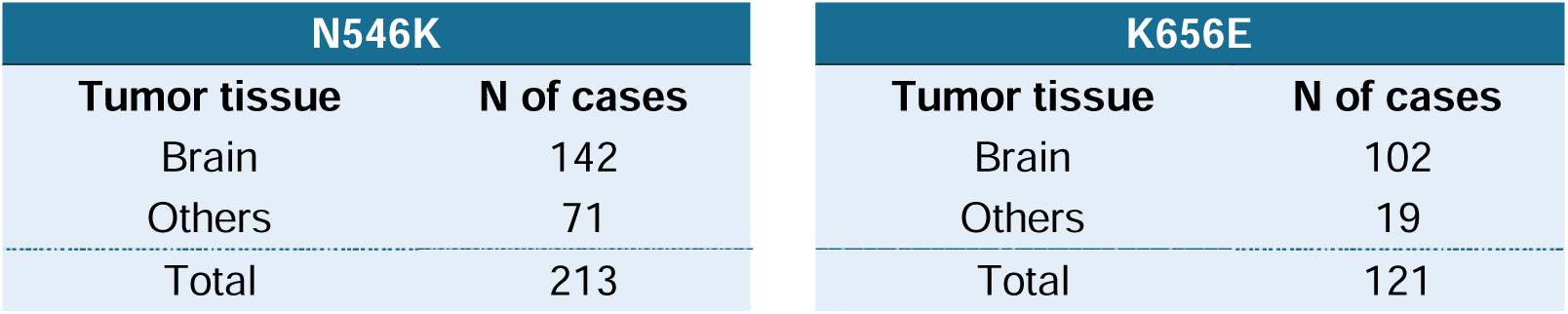
Number of cases in the GENIE database harboring the N546K or K656E mutation. Samples are categorized based on their classification as brain tumors or other tumor types.

**Table 2.** Number of patients with *FGFR1*-mutated brain tumors in the GENIE database. Samples are distributed based on whether they present only one of the hotspots mutations N546K or K656E (“Single”) in *FGFR1* or they display additional missense mutations in *FGFR1* (“Double/Multiple”).

| N546K |  | K656E |  |
| --- | --- | --- | --- |
| *FGFR1* hits | N of cases | *FGFR1* hits | N of cases |
| Single (hotspot only) | 126 | Single (hotspot only) | 56 |
| Double/Multiple | 16 | Double/Multiple | 46 |
| Total | 142 | Total | 102 |

**Table 3.** Number of *FGFR1*-mutated brain tumor samples in the GENIE cohort, categorized by tumor grade and divided in Single and Double/Multiple (same as Table 2). Low-grade: tumor types with grade 1 or 2; high-grade: tumor types with grade 3 or 4; unknown: tumor types that could not be assigned.

| N546K |  |  |  |
| --- | --- | --- | --- |
| Tumor grade | *FGFR1* hits |  |  |
|  | Single (hotspot only) | Double/Multiple | Total |
| Low-grade | 52 | 10 | 62 |
| High-grade | 65 | 5 | 70 |
| Unknown | 17 | 1 | 18 |
| Total | 134 | 16 | 150 |

**Table 3.** Number of *FGFR1*-mutated brain tumor samples in the GENIE cohort, categorized by tumor grade and divided in Single and Double/Multiple (same as Table 2).
| K656E |  |  |  |
| --- | --- | --- | --- |
| Tumor grade | *FGFR1* hits |  |  |
|  | Single (hotspot only) | Double/Multiple | Total |
| Low-grade | 31 | 31 | 62 |
| High-grade | 20 | 14 | 34 |
| Unknown | 5 | 2 | 7 |
| Total | 56 | 47 | 103 |

### Proximal interactome of wild-type and mutant FGFR1 receptors

To investigate how different mutations rewire the network of molecular interactions engaged by the FGFR1 receptor, we employed a systematic interactome-profiling approach using the BioID strategy coupled to mass spectrometry (MS). The approach is based on the biotinylation of proteins within a 10 nm radius of the bait mediated by the mutant enzyme BirA* in the presence of biotin (Roux *et al*, 2012) and allows the identification of interactions hard to capture with standard co-immunoprecipitation strategies. For the purpose of this study, we chose to focus on the two hotspots N546K and K656E, the germline variant R661P, previously identified by our research team (Rivera *et al*., 2016), and both combinations *in cis* (N546K/R661P and K656E/R661P) as models of “hotspot + secondary” double mutants (Fig. 2A-B). Toward this aim, the Flp-In T-REx HEK293 system has been used to generate inducible, stable cell lines expressing the coding sequences of WT and mutant FGFR1 receptors, fused to the mutant enzyme BirA* and Flag (Fig. 2B). The expression and activation (phosphorylation of activation loop Tyrosine residues 653/654) of the six FGFR1-BirA*-Flag fusion proteins were validated by western blot (Fig. 2C), demonstrating proper kinase function of the BirA*-fused C-terminal domain.

**Figure 2.**
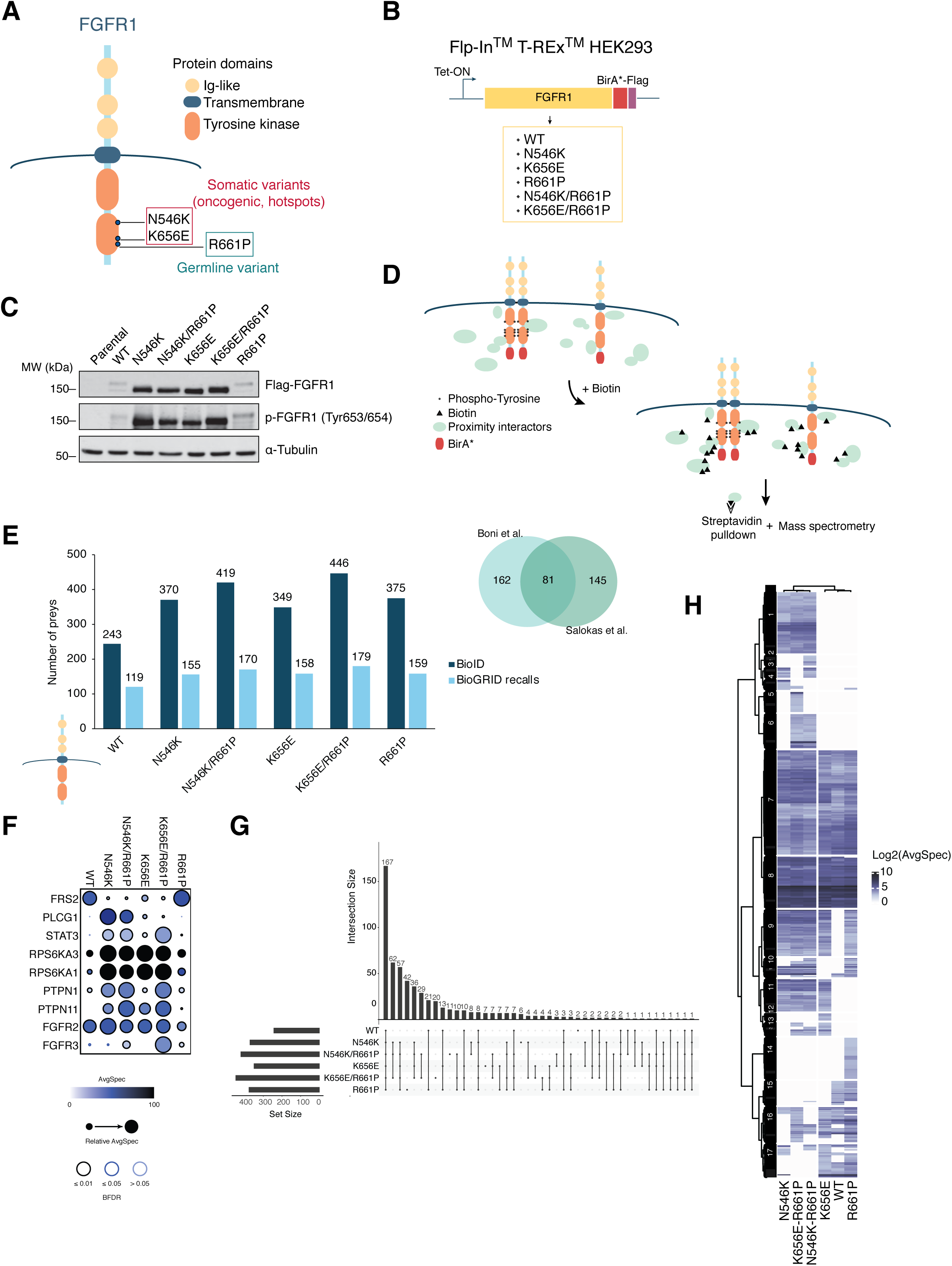
Proximal interactome profiling (BioID-MS) of WT and mutant FGFR1 proteins. **A)** Scheme of the FGFR1 receptor representing functional domains and SNVs investigated in the present study: the hotspots N546K, K656E and the germline variant R661P. **B)** Stable inducible HEK293 (Flp-In T-REx) cell lines have been generated to express the coding sequence of WT and the 5 mutant FGFR1 fused mutant BirA* enzyme and Flag. **C)** Western blot of the parental (empty-Flag) and the six Flp-In T-REx HEK293 cell lines used in the present study, showing expression and activation (phosphorylation of Tyr653/654) of FGFR1, after 24h of Tet induction. **D)** A schematic outline of the BioID-MS screen: in the presence of biotin the BirA* enzyme fused to both active (phosphorylated dimers) and inactive FGFR1 molecules catalyzes the biotinylation of proximal interactors, which are isolated through streptavidin-mediated pull-down and analyzed through MS. **E)** Number of identified preys (after filtering) as well as BioGRID recalls for each bait are represented. Right: numbers of shared and non-shared interactors of WT FGFR1 between our list and the one published by (Salokas *et al*., 2022). **F)** Dot plot highlighting FGFR1 *bona fide* interactors and their abundances (AvgSpec) among the different baits. **G)** Upset plot showing numbers of proximal interactors among different combinations of intersections of the different baits. **H)** Heatmap of an unsupervised hierarchical clustering analysis using bait abundances (Log2AvgSpec). 17 distinct clusters have been identified. AvgSpec = Average of spectral counts.

The rationale of our experimental design was based on a BirA* enzyme fused to the C-terminus of the FGFR1 protein, providing labelling of intracellular factors in the proximity of the receptor, which have been purified through streptavidin affinity pull-down and identified through mass-spectrometry (Fig. 2D). Biotinylation of proximal interactors was confirmed for all the baits by western blotting (Appendix Fig. S2). Despite the observed differences in protein amounts among conditions (Fig. 2C, discussed in the next section) we did not obtain any bias in the outcome of our interactome profiling, as indicated by comparable total numbers of identified unique preys (Fig. 2E, Dataset EV1). Our BioID capture was able to recall FGFR1 (WT) interactors reported in the BioGRID database (Oughtred *et al*, 2021), including proximal interactors (n = 81) identified in a recent comprehensive study using human RTKs as baits expressed through the same cell model (Flp-In T-REx HEK293) (Salokas *et al*, 2022) (Fig. 2E, Dataset EV1). This data supports the reliability of our experimental procedures and, on the other hand, highlights potential novel, so far unreported FGFR1 protein partners. Among the recalled preys, we also retrieved many well established FGFR1 *bona fide* interactors, including other members of the FGFR family (FGFR2 and FGFR3), modulators as protein tyrosine phosphatase non-receptor type 1, 11 (PTPN1, PTPN11) and downstream signaling effectors as fibroblast receptor substrate 2 (FRS2), phospholipase C gamma 1 (PLCγ/PLCG1) and signal transducer and activator of transcription 3 (STAT3) (Fig. 2F). The composition of each interactome was compared, showing both unique and shared preys among FGFR1 mutants and with the WT protein (Fig. 2G). The most abundant group (n = 167) was formed by proteins detected in every condition, confirming that differences in protein expression had not a strong impact in the BioID-MS and highlighting a core of functionality not affected by the mutations considered in this study (Fig. 2G). To further elucidate similarities and divergences among the WT and mutant proteins, we performed an unsupervised hierarchical clustering analysis. The germline R661P clustered differently (and closer to the WT protein), compared to the hotspot mutants, both single and double (Fig. 2H). On the other hand, although they clustered similarly to their respective single mutants N546K and K656E, double mutants displayed specific interactors (Fig. 2H, Clusters 2, 5 and 6; Dataset EV1). This overview of the interactome already indicated the uniqueness of the R661P variant and potential modulatory effects that it may exert when combined with the oncogenic mutations.

### FGFR1 oncogenic mutants present higher levels of protein accumulation

According to their activating nature, we expected the two hotspot mutations N546K and K656E to appear with increased phospho-FGFR1 (p-FGFR1) levels in western blot compared to the WT protein. We did observe stronger intensity p-FGFR1 bands for the two single mutants, while the double mutants mirrored this pattern, without showing any clear change (Fig. 2C). However, by looking at the total Flag-FGFR1 levels, we observed a similar trend, implying that the enhanced signal was not solely due to intrinsic phosphorylation, but also reflected increased protein accumulation (Fig. 2C). To further test this, total Flag-FGFR1 protein levels from multiple independent experiments were quantified, revealing a significant and high (4-6-fold change) increase in the levels of accumulation of the oncogenic mutant proteins, both in single and double settings (Fig. 3A). On the contrary, when phopsho-FGFR1 levels were quantified and corrected for total Flag-FGFR1 levels, the differences between oncogenic mutants and WT protein were less pronounced and statistically non-significant, except for N546K (Fig. 3A). Similar total and phosphorylated protein profiles were observed in cells previously serum-starved, indicating that, in our settings, differences among conditions do not depend on the presence of growth factors (Fig. EV3A). Notably, when we looked at specific interactors in BioID for this group (Fig. 2H, Cluster 11), we found an enrichment in GO categories related to protein transport and vesicles trafficking, which might indicate altered regulation of receptor turnover (Fig. 3B), potentially affecting recycling and degradation rates (Miaczynska, 2013). Consistent with previous findings on N546K (Lew *et al*., 2009), HEK293 cells overexpressing an oncogenic mutant FGFR1 (N546K and K656E, but also double mutants N546K/R661P and K656E/R661P) displayed a transformed phenotype with changes in morphology toward a stem-like shape and higher tendency to detach from the plate (Fig. EV3B), large nuclei and reduced cytoplasmic volume (Fig. 3C). In addition, we observed different FGFR1 subcellular localization between WT and oncogenic protein-expressing cells. While WT and R661P mutant display classic receptor localization, mainly in the plasma membrane with little cytoplasmatic involvement, oncogenic mutants accumulate at high levels in intracellular compartments (Fig. 3C), reinforcing the hypothesis of a dysregulated receptor transport for these conditions. Coherent with these changes in localization, analysis of groups of interactors identified exclusively for R661P and WT FGFR1 (Fig. 2H, Clusters 14-15; Dataset EV1) were found enriched in plasma membrane and cytoskeleton-associated proteins (Fig. EV3C).

**Figure 3.**
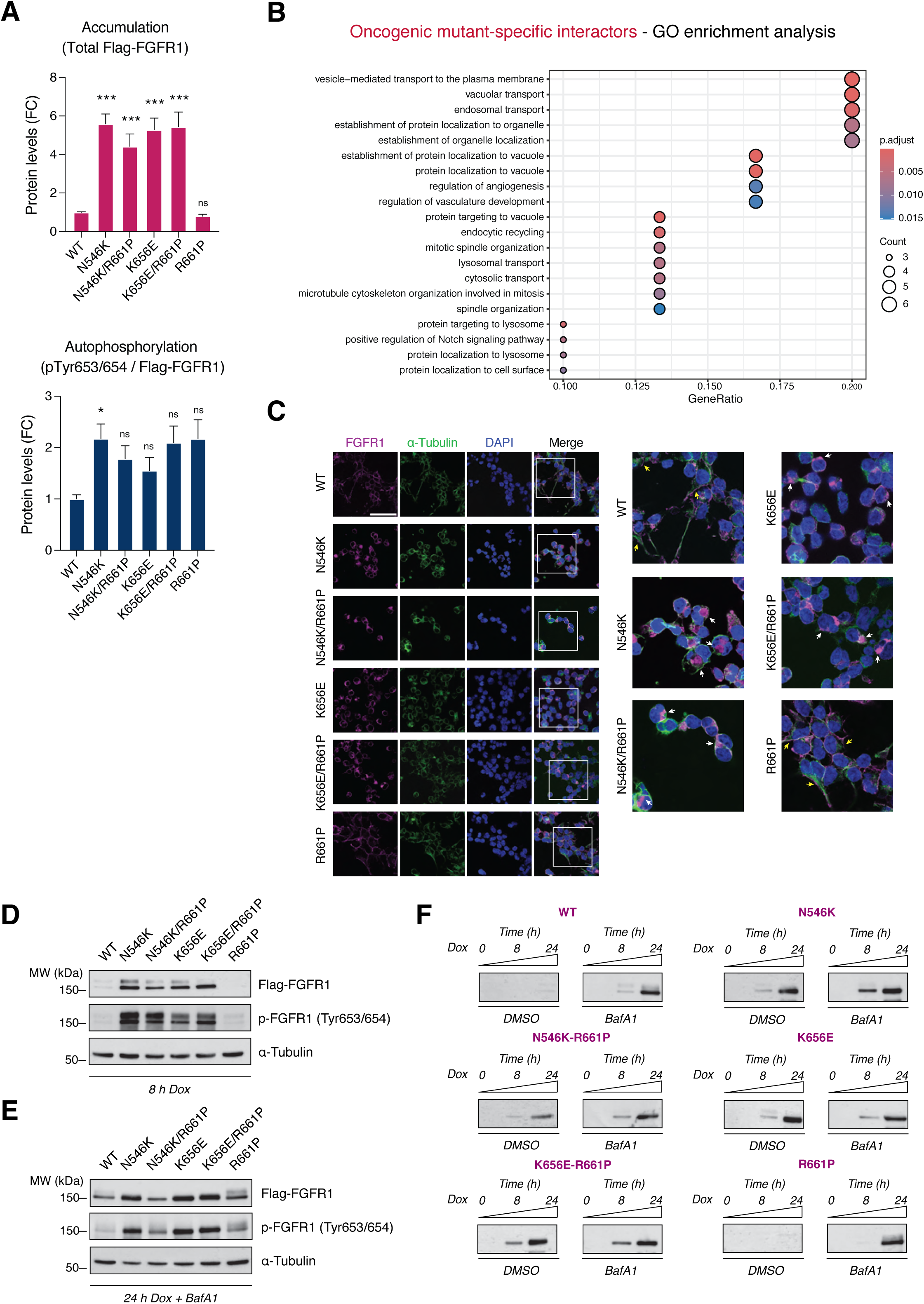
N546K and K656E mutated receptors accumulate at higher levels in HEK293 cells. **A)** Bar plot of relative amounts (3 technical quantifications of western blots from n = 6 independent experiments) of total Flag-FGFR1 and ratio phospho-Tyr653/654 / Flag-FGFR1 of WT and mutant FGFR1 proteins (as in Fig. 2C) obtained using the six T-REx HEK293 cell lines, representing differences in levels of protein accumulation and intrinsic autophosphorylation, respectively. Fold change (FC) values, obtained normalizing against WT levels, are indicated. Error bars represent standard errors of the mean ± SEM values. Significant comparisons against WT values are indicated (Krustal-Wallis test, p value * ≤ 0.05, ** ≤ 0.01, *** ≤ 0.001). **B)** Dot plot of GO enrichment analysis using the list of genes included in Cluster 11 of the heatmap in Fig. 2H (shared interactors among the 4 hotspot-expressing cell lines). The size size and color of dots represent numbers of matched genes and adjusted p-value (p.adjust), respectively. **C)** Immunofluorescence staining of FGFR1 protein (pink) and α-Tubulin (green) using the T-REx HEK293 FGFR1 cell lines, displaying differences in cellular localization of FGFR1 receptors. Right panels: zoom in (white squares in left panels). Arrows highlight different patterns of FGFR protein in the oncogenic mutants (white arrows indicating accumulation in intracellular compartments), compared with WT and R661P (yellow arrows indicating FGFR1 localization to the plasma membrane). Scale bar: 50 µm. **D-E)** Western blot assays revealing Flag-FGFR1 expression and tyrosine phosphorylation after 8 hours (D) and 24 hours (E) of Tet-induction. In (E) cells have been treated in parallel with the lysosome inhibitor Bafilomycin A1 (200 nM). **F)** Time course of Flag-FGFR1 protein synthesis showing differential accumulation rates and patterns among the different mutants and in presence or absence of BafA1. Equal amounts of loaded protein have been assessed through Ponceau staining (Appendix Fig. S3). GO, Gene Ontology.

Considering the homogenous expression ensured by the Flp-In system (transcript levels are represented in Fig. EV3D, with no condition exceeding a two-fold change variation), the increased levels observed at 24h post-Tet induction are most likely due to post-translational mechanisms occurring during this time frame. To explore early events occurring after FGFR1 mutant synthesis and accumulation, we collected cell lysates at 8 hours of induced expression. At 8 hours, all mutants, including hotspots, revealed a WT-like profile, with two distinct bands, highlighting similarities in the post-translational processes they undergo at early stages (Fig. 3D). However, phosphorylation of oncogenic mutant receptor is already enhanced at this step, confirming that hyperphosphorylation is an early event driven by the amino acid changes N546K and K656E, with higher power exerted by the first one. To determine whether these mutations are leading to degradation escape, we treated the cells with Bafilomycin A1 (BafA1), a potent inhibitor of vacuolar H^+^ ATPases and therefore of lysosomal function (Yoshimori *et al*, 1991). Upon BafA1 addition, WT and R661P FGFR1 displayed an “oncogenic mutant-like” pattern with the lower molecular weight band that accumulates over the upper one (Fig. 3E), suggesting that, in normal conditions, i) this low-molecular weight protein form is the most frequently degraded in lysosomes in WT settings and ii) oncogenic mutant FGFR1 receptors escape lysosomal degradation, accumulating only the faster-resolving band (Fig. 2C and Fig. 3E). These findings are further substantiated in Fig. 3F, where the blots show the time course of expression of WT and mutant FGFR1 proteins at 0, 8 and 24h post Tet-induction, in the presence or absence of BafA1.

### R661P mutation partially rescues lysosomal degradation of oncogenic proteins

To further investigate whether oncogenic mutants evade vesicular degradation pathways, we induced receptor internalization and lysosomal degradation by stimulating the FGFR1-expressing HEK293 cells with the FGFR1 ligand FGF2 (Fig. 4A). In parallel, to avoid accumulation of newly synthesized protein, we inhibited mRNA translation by treating the cells with cycloheximide (CHX) (Fig. 4A). While WT levels decreased over time, the oncogenic mutant K656E escaped degradation, maintaining stable protein levels (Fig. 4B-C). Notably, this degradation was partially rescued in the double mutant (Fig. 4B-C), with protein levels decreasing to 50% at 6 hours, indicating a novel modulatory mechanism played by the R661P variant over the oncogenic mutant K656E. Supporting these results, upon treatment with Bafilomycin, degradation was prevented and similar levels of WT and K656E/R661P proteins (38 and 31%, respectively) were recovered at 6 hours (Fig. EV4A-B). The destabilization exerted by R661P was confirmed for the N546K oncogenic mutant, as the N546K/R661P double mutant also rescued partial degradation (45% of starting protein levels) (Fig. 4D-E). These results highlight similar mechanisms wielded by the R661P to enhance degradation of both oncogenic mutants.

**Figure 4.**
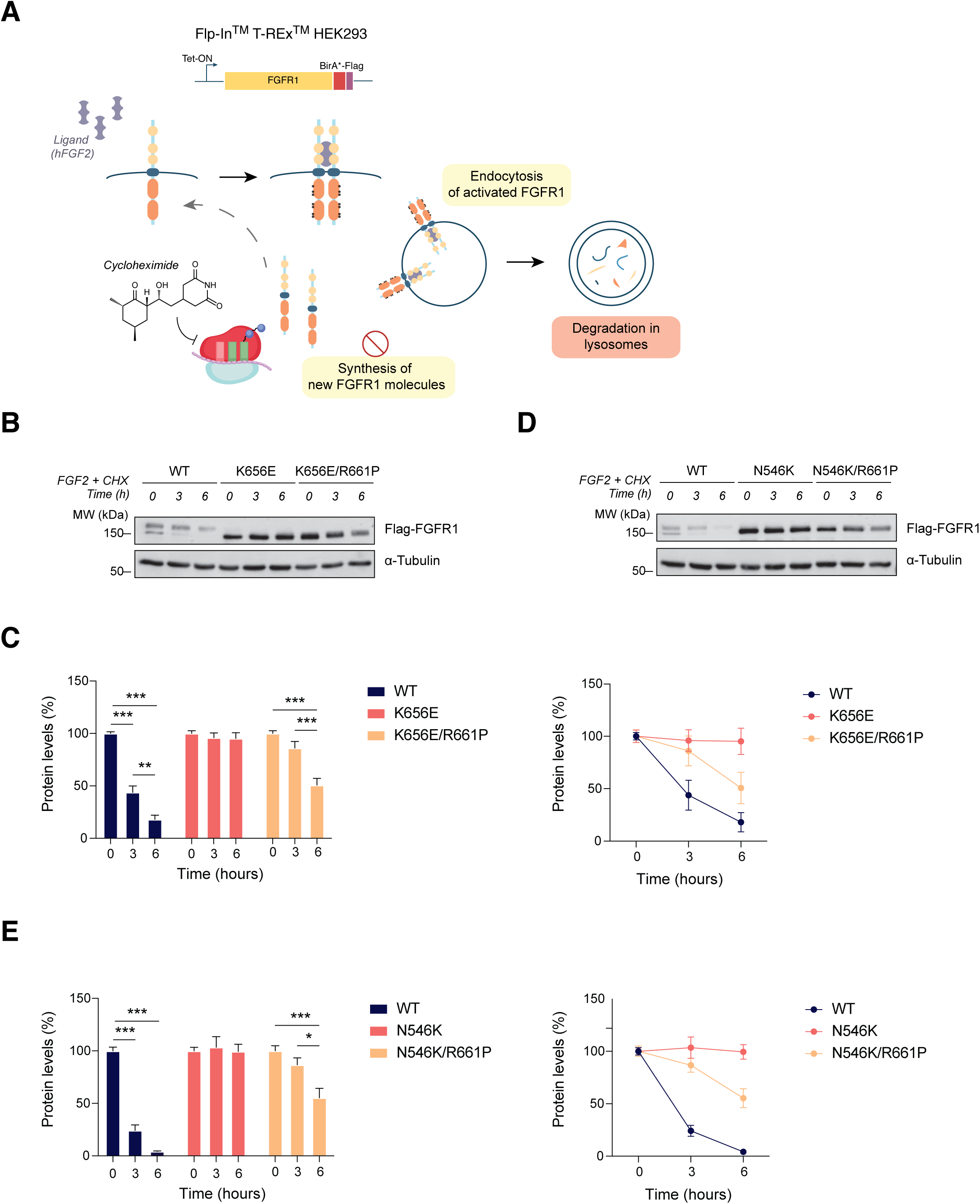
R661P reduces K656E and N546K mutant protein stability by rescuing lysosomal degradation. **A)** Receptor stability experimental design: HEK293 cells expressing WT and mutant FGFR1 are stimulated with human FGF2 (hFGF2, 25 ng/mL) to induce receptor internalization and degradation and, in parallel, with cycloheximide (100 µg/mL) to block protein synthesis. **B and D)** Western blot analysis of a representative time-course experiment comparing WT FGFR1, K656E (B) and N546K (D) single and double mutants showing total Flag-FGFR1 protein levels at 0, 3 and 6 hours post-treatment with FGF2 and cycloheximide (CHX). **C and E)** Relative amounts of WT, single and double mutant FGFR1 protein obtained by quantifying western blots from n = 3 independent experiments. Values have been normalized (FC) against each relative 0h reference values. Data is represented by mean ± SEM and significant variations in protein amounts against each specific reference values have been indicated (ANOVA test, p value * ≤ 0.05, ** ≤ 0.01, *** ≤ 0.001) in the bar plots.

### The R661P variant destabilizes PLCγ binding acquired by N546K mutant

As mentioned above, the proximity interactome revealed differences in the abundance of crucial downstream signaling mediators between WT FGFR1 and oncogenic mutant baits (Fig. 2F). Among those, we decided to further investigate the interaction with PLCγ since it was found strongly increased in both N546K and N546K/R661P captures, pointing to extended binding and activation of downstream signaling (Fig. 5A). PLCγ is encoded by the *PLCG1* gene and, upon binding to RTK receptors is activated and mediates the cleavage of hydrolyzed phosphatidylinositol-4, 5-bisphosphate (PIP2) to produce inositol-1,4,5-triphosphate (IP3) and diacylglycerol (DAG), which in turn regulate multiple cellular processes in a tissue-specific manner (Yang *et al*, 2012). To further understand this interaction at the molecular level, we leveraged the structure prediction algorithm Alphafold3 (AF3) (Abramson *et al*, 2024) to generate a 3D model of the FGFR1-PLCγ complex (Fig. 5B) and confirmed that the model correctly recapitulates experimentally validated interactions between specific domains (Hajicek *et al*, 2019) and superposes with the crystalized structure of the FGFR1 active kinase domain (Appendix Fig. S4). In line with our interactome results, the residue Asn546 (contrary to Lys656 and Arg661) is located at the surface of interaction between FGFR1 and PLCγ (Fig. 5B), and the side chain of the mutated residue (lysine) is exposed toward the interface of interaction, suggesting that it might be stabilizing the binding (Fig. 5B). Despite this interaction was also strong in the double mutant N546K/R661P, it appeared attenuated (Fig. 5A), potentially indicating reduced affinity compared to the respective single mutant. The increased ability of the N546K single-and double-mutant protein to bind and phosphorylate PLCγ1 was validated through co-immunoprecipitation experiments (Fig. 5C). Once again, the presence of the R661P mutation was found to weaken this interaction and consequent activation of PLCγ, with lower levels of immunoprecipitated phosphorylated proteins, highlighting another modulation mechanism played by R661P on the activating potential of the N546K mutation.

**Figure 5.**
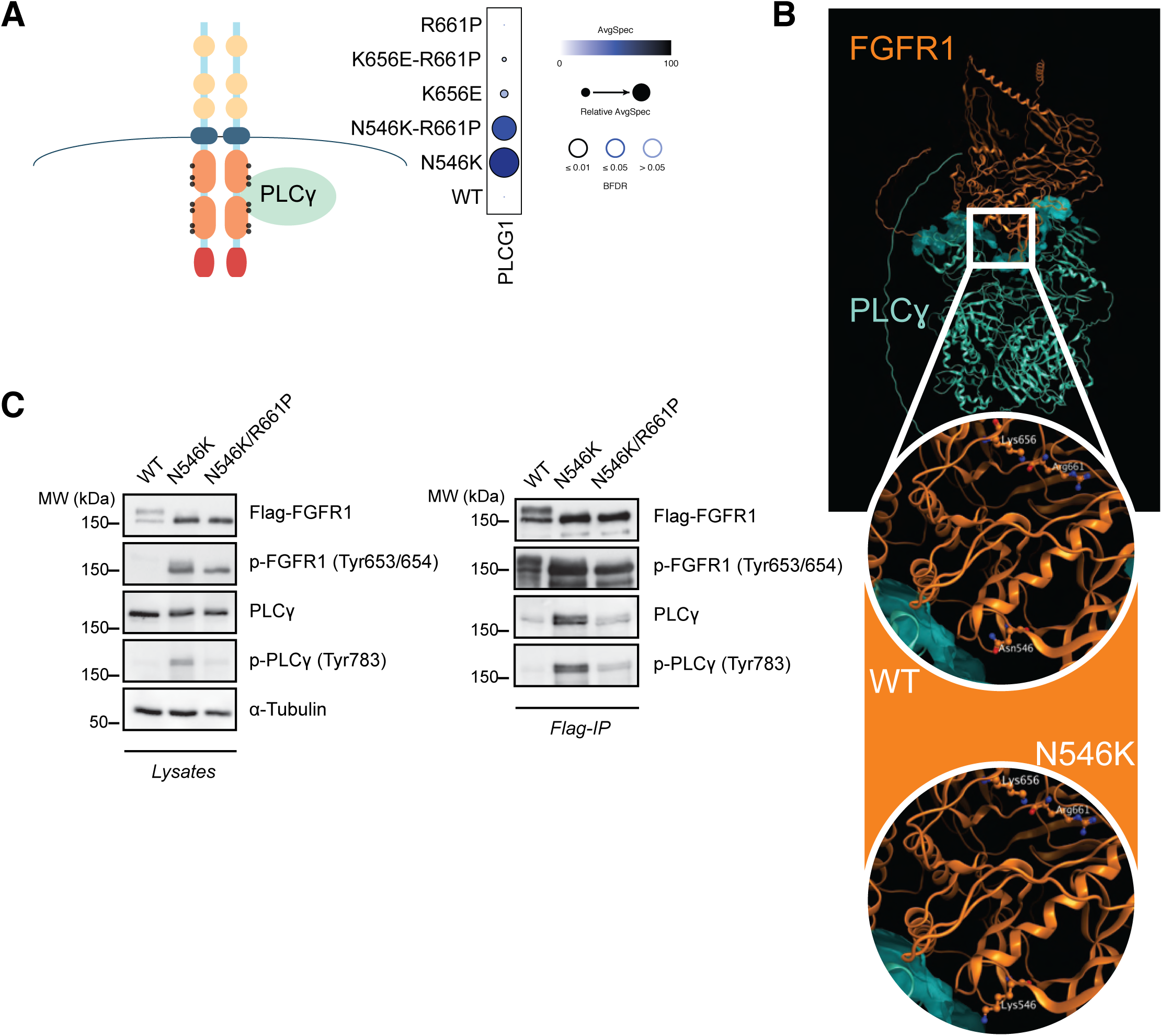
N546K/R661P double mutant interacts with PLC**γ** with reduced affinity compared to N546K. **A)** Schematic representation of activated FGFR1 bound to PLCγ (left) and dot plot with relative abundances (AvgSpec) of PLCγ protein for WT and mutant FGFR1 obtained through the BioID-MS screening. **B)** Predicted 3D structure of FGFR1-PLCγ complex obtained using AF3. The square highlights a zoomed area with the 546 residue in WT (Asn) and mutant (Lys) FGFR1 protein, locating in the interface of FGFR1-PLCγ interaction. The residues Lys656 and Arg661 are also highlighted, to show how residue 546 is the only one close to the interface of interaction and supporting a role for the N546K mutation in enhancing the binding between the two proteins. **C)** Western blot of total lysates and the co-immunoprecipitation (IP) of total and phosphorylated FGFR1 and PLCγ proteins, using the T-REx HEK293 stable cell lines. AvgSpec = Average of spectral counts.

### R661P severely impairs the proliferative potential of oligodendroglioma cells

It has been proposed that DNET and other types of glioneuronal tumors originate from oligodendrocyte lineages and express molecular signatures of oligodendrocyte precursors (Duan *et al*, 2024; Luzzi *et al*, 2019; Matsumura *et al*, 2013). To study the impact of the mutations investigated so far on a model that could better recapitulate the molecular background of *FGFR1*-mutated brain tumors we used the human oligodendroglioma cell line (HOG) and we applied CRISPR/Cas9 to generate isogenic FGFR1-mutant HOG cell lines (Fig. 6A). HOG cells have been shown to recapitulate immature oligodendrocyte lineage and have been used in previous reports to study oligodendrocyte function (Buntinx *et al*, 2003; De Kleijn *et al*, 2019; Post & Dawson, 1992). They are hemizygous for *FGFR1*, as they harbor a genomic deletion in chr8p.23 encompassing the *FGFR1* locus (Fig. 6A). This aspect was crucial to study *FGFR1* mutations without WT gene expression, as it allowed us to force our cellular systems to reveal variant-associated phenotypes. Thus, two independent clones for the same *FGFR1* mutations characterized in the interactome profiling (N546K, K656E, R661P, N546K/R661P and K656E/R661P) were isolated and correct editing was confirmed (Appendix Fig. S5). Mutant clones presented no apparent differences in morphology except from R661P-expressing cells, which tend to form more compacted clusters, with cells establishing more surface contacts (Fig. EV6A). Molecularly, this could be explained by a different polarization mediated by FGFR1-R661P at the plasma membrane, engaging interactions with different actin-interacting proteins than the WT protein, as indicated by analysis of the R661P-specific BioID cluster 14 (Fig. EV3B, Dataset EV1). Strikingly, colony formation assays pointed to the R661P germline variant as the mutation with the most severe effect, almost completely abolishing the clonogenic potential of HOG cells (Fig. 6B). In these assays, both double mutants displayed an intermediate phenotype between the one caused by the germline variant and their respective oncogenic single mutants (Fig. 6B).

**Figure 6.**
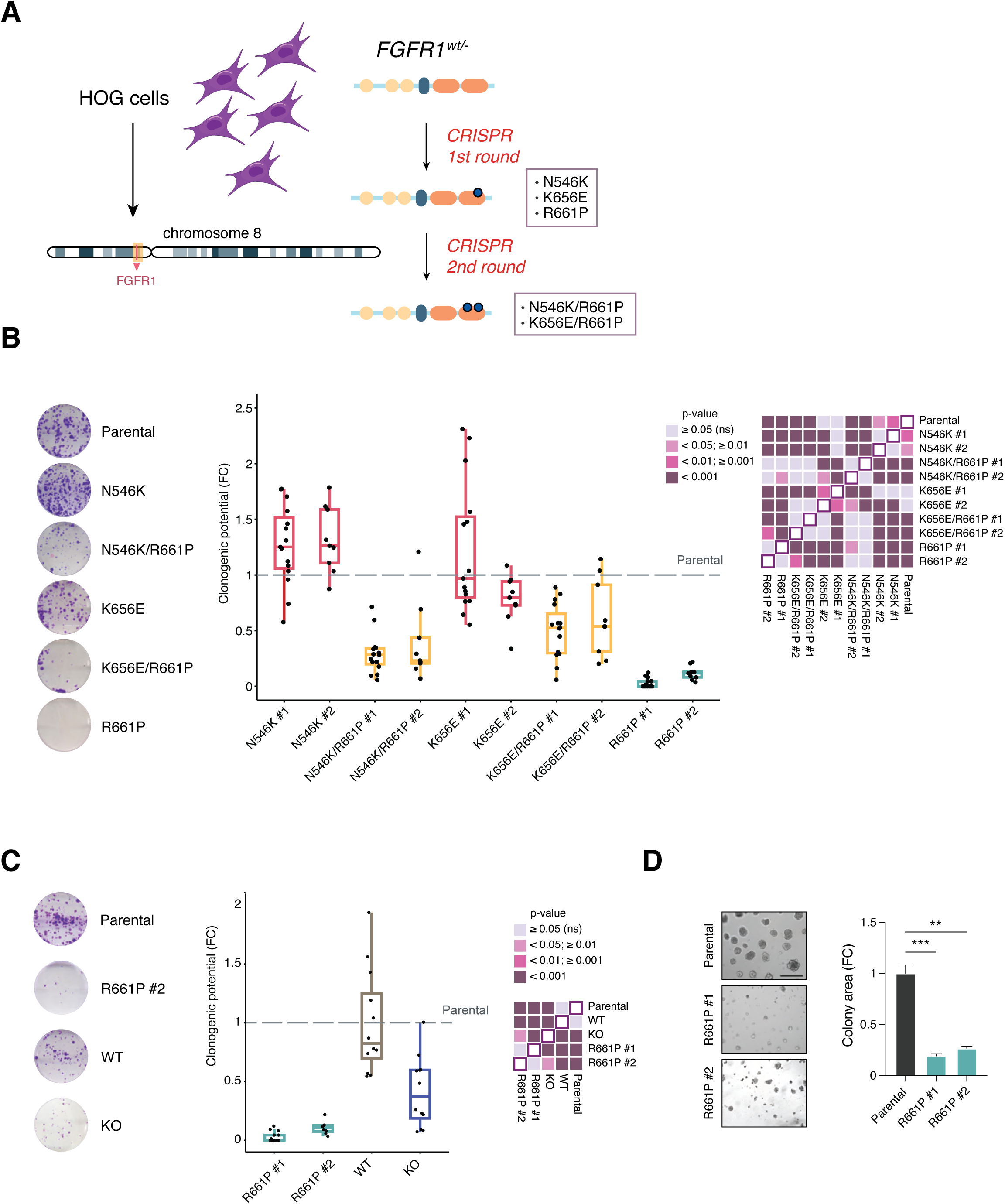
Defects in self-renewal and changes in phenotype imposed by R661P mutation. **A)** Schematic representation of the CRISPR design to generate FGFR1 single-and double-mutant clones using the HOG cell line. Genomic deletion in *FGFR1* locus in chromosome 8 (chr.8p11.23) identified in these cells has been represented**. B)** Colony forming assays comparing CRISPR-edited, FGFR1-mutant HOG clones. Both clones for each mutation (N546K, K656E, N546K/R661P, K656E/R661P and R661P) have been assayed. Microscope captures of representative wells of clones #1 for each mutant have been included on the left and quantification of n ≥ 3. experiments have been plotted, normalized against the parental cell line (dashed line). Replicates are represented by black dots. Results of statistical analysis of multiple comparisons (ANOVA test) have been plotted in the heatmap on the right. **C)** Colony forming assays comparing the two R661P clones with an WT clone and a KO clone. Representative wells of the second R661P clone, WT and KO clones are indicated. Quantification of n ≥ 3. experiments have been plotted, normalized against the parental cell line (dashed line). Results of statistical analysis of multiple comparisons (ANOVA test) have been plotted in the heatmap on the left. **D)** Matrigel 3D colony forming assays comparing the two R661P clones with the parental cell line. Captures of a representative field displaying colonies formed at experimental endpoint (day 7) are reported on the left. Scale bar: 200 µm. On the right: bar plot of quantified colony areas, normalized against control condition (Parental cell line). Mean ± SEM values (n ≥ 3 independent experiments) and significant differences have been indicated (Krustal-Wallis test, p value * ≤ 0.05, ** ≤ 0.01, *** ≤ 0.001).

To study whether the R661P proliferative defect was associated with a loss of FGFR1 function, we also generated a knock-out (KO) clone (Fig. EV6C). Although *FGFR1*-KO cells showed impaired self-renewal ability compared to the parental cell line, this effect was less pronounced than for R661P-mutant cells (Fig. 6C). This observation suggests that the phenotypic consequences of the R661P variant are probably linked to a switch, rather than a loss, of receptor function, which leads to more dramatic effects in HOG cells compared to total loss of *FGFR1* expression. Finally, we tested a FGFR1 WT-clone isolated during CRISPR experiments, which exhibited no significant differences with the parental cell line (Fig. 6C). This result provides additional evidence that the observed phenotype was mutation-specific, and not a stochastic fluctuation of proliferation rates in HOG cell population. Defective self-renewal ability of the two R661P clones was confirmed in 3D-Matrigel colony formation assays, with a reduction in the size of colony area of 70-80% (Fig. 6D). In summary, these results unveiled severe defects in the proliferative potential of oligodendroglioma cells caused by the R661P mutation.

### R661P-mutant oligodendroglioma cells present changes in developmental gene expression, recovered in double mutant cells

To investigate the molecular events driving the phenotypic changes observed above in HOG cells, we performed differential gene expression analysis through RNAseq using the two R661P mutant HOG clones and the parental cell line, identifying n = 338 of differentially expressed (DE) genes shared by the two clones, representing the 22% and 26% of total DE genes for clone #1 and clone #2, respectively (Fig. 7A, Dataset EV2). Gene ontology analysis of DE genes in FGFR1-R661P HOG cells revealed enrichment for development-related categories, some of them specific for brain tissue formation and differentiation (Neural Crest Cell development, Axon guidance, Axonogenesis), alongside with cellular processes (Epithelial cell proliferation, Negative regulation of cell adhesion and Cell-matrix adhesion) which may underlie the R661P cell phenotype discussed in the previous section (Fig. 7B). We selected key genes belonging to these categories that were found dysregulated in R661P clones and we validated them through RT-qPCR (Fig. 7C). These genes encode for described master transcriptional regulators (*SOX9, GLI2, HIF1A*), signaling factors (*WNT5A, NRP1, SEMA3E*), and surface glycoproteins (*THBS1, FN1*) with central roles during development, neuro-glial cell fate decisions and oligodendrocyte differentiation (Allan *et al*, 2021; Blake *et al*, 2008; Endo & Minami, 2018; Finzsch *et al*, 2008; Hassel *et al*, 2023; Oh & Gu, 2013; Qi *et al*, 2003; Sherafat *et al*, 2021).

**Figure 7.**
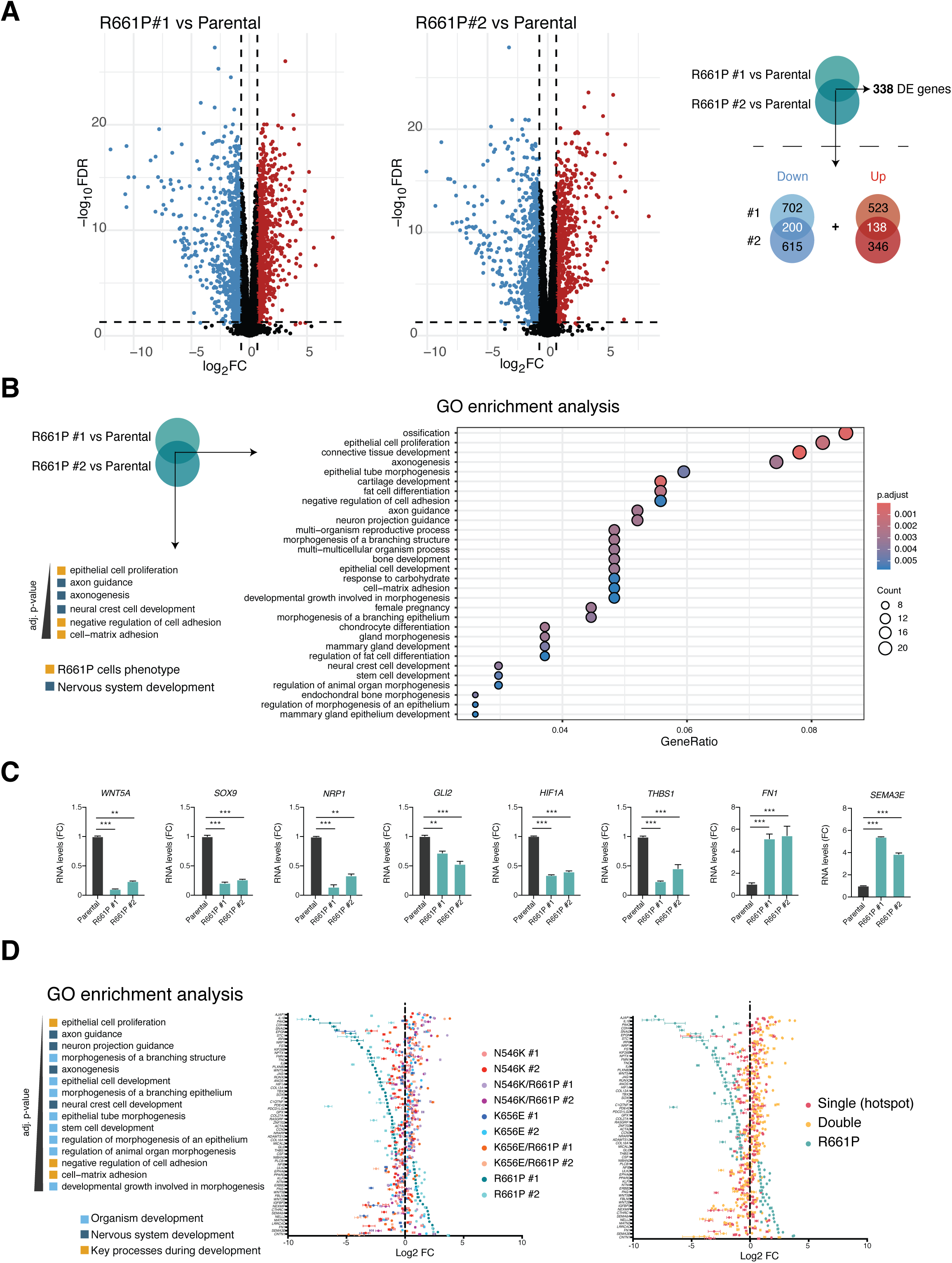
R661P mutant HOG cells show dysregulation of development and neurodevelopment-associated genes, rescued in double mutants. **A)** Volcano plots representing downregulated (red dots) and upregulated (blue dots) genes differentially expressed genes comparing each R661P clone with the Parental cell line. Numbers of specific and common DE genes are specified in Venn diagrams (right). **B)** Dot plot of GO enrichment analysis of R661P vs Parental differentially expressed (DE) genes, using common DE genes obtained from both comparisons showed in A) as input list. Significantly enriched (adj. p value ≤ 0.05) have been represented. Size and color of dots represent number of matched genes and adj. p value, respectively. **C)** Selected GO categories for further analysis are illustrated. RT-qPCR validation of key genes, comparing RNA levels in R66P clones vs Parental HOG cells. Mean ± SEM values from n =2 independent experiments have been plotted and significant differences have been indicated (ANOVA test, p value * ≤ 0.05, ** ≤ 0.01, *** ≤ 0.001). **E)** Dot plots representing Log_2_FC values of RNAseq DE analysis obtained comparing each FGFR1 mutant clone with the Parental cell line. Error bars represent SD among triplicates. Selected genes (n= 64) that have been plotted are unique genes identified in the GO categories listed on the left (selected from GO in B). Left plot: each independent clone has been represented with a different color. Right plot: Single mutant clones (hotspots N546K #1 and #, K656E #1 and #2) have been represented in red; double mutant clones (N546K/R661P #1 and #2, K656E/R661P #1 and #2) have been represented in light orange and R661P #1 and #2 clones have been represented in blue. DE = Differentially expressed

To assess whether the mentioned processes are also altered in other FGFR1 mutants, or they are R661P-specific, we also performed RNAseq using N546K, K656E and double mutants N546K/R661P and K656E/R661P clones. By extending the number of categories of interest obtained in Fig. 7B to more general development-related categories, we plotted differential gene expression values obtained by comparing each mutated clone with the parental cell line, focusing on unique genes belonging to these groups (Fig. 7D). Despite the presence of the R661P mutation, double mutant N546K/R661P and K656E/R661P HOG cells showed a regression toward parental expression levels, with a profile close to the one traced by N546K and K656E clones (Fig 7D). These observations indicate that the presence of the hotspot *in cis* is able to compensate for potential defects caused by the germline allele and point to a novel model of susceptibility for the *FGFR1* allele R661P.

## Discussion

Multiple mutations *in cis* in tumor driver genes are rare events, mostly associated with recurrence and acquired therapy-resistant phenotypes (Barber *et al*, 2013; Castaneda-Gonzalez *et al*, 2022; Kondrashova *et al*, 2017; Stewart *et al*, 2015). By interrogating GENIE project data, the largest tumor cohort publicly available so far, we confirmed the recurrence of secondary variants along with K656E and N546K hotspots in brain tumors, more frequently found with the first one. Next, we employed complementary models and strategies, providing multiple evidence on both molecular mechanisms and phenotypes triggered by double mutant FGFR1 receptors. FGFR1 and, in general, RTK signaling is spatially and temporally controlled by fine-regulated intracellular vesicular trafficking (Goh & Sorkin, 2013; Hinsch *et al*, 2023; Miaczynska, 2013). The data presented in this study highlights an oncogenic mechanism for the two hotspot mutations that relies on altered protein transport and evasion of degradation processes, resulting in the accumulation of active (phosphorylated) FGFR1 receptor in cells rather than solely on increased enzymatic activity, as proposed in previous works (Bennett *et al*., 2016). Nevertheless, mutant FGFR1 N546K also shows significant increased autophosphorylation (Fig. 3A), in line with a described biochemical model that proposed a faster and ligand-independent autophosphorylation (Lew *et al*., 2009), which could mediate the activation of downstream signaling pathway and, in parallel, escaping of degradation mechanisms. Our proximal interactome profiling identified enhanced binding to PLCγ for the N546K mutant; one possibility is that the stronger interaction prevents FGFR1-N546K protein to be targeted for degradation. This hypothesis is supported by the results obtained with the double mutant, which loses affinity for PLCγ and partially rescues degradation.

Little experimental evidence is available in literature for K656E activating molecular features (Fisher *et al*, 2021; Hart *et al*., 2000). However, the analogous variant in FGFR3 (K650E) in germline carriers causes skeletal dysplasia syndromes (Foldynova-Trantirkova *et al*, 2012; Wilcox *et al*, 1998), and has been reported to enhance autophosphorylation and induce proliferative phenotypes in cells (Bellus *et al*, 2000; Monsonego-Ornan *et al*, 2002; Naski *et al*, 1996). Notably, ectopic expression of FGFR3 K650E resulted in a similar loss of the upper band in western blot experiments, indicating analogous changes in the combination of post-translational modifications induced by the amino acid change (Bellus *et al*., 2000; Monsonego-Ornan *et al*., 2002). In fact, FGFR1 Lys656 was included in a screening of multiple FGFR1 lysine residues mutants, which resulted in escaped lysosomal degradation (Haugsten *et al*, 2008), reinforcing our premise that oncogenic mutants affecting this conserved residue act through dysregulation of degradation pathways in FGF receptors.

For pediatric and epileptogenic low-grade glioneuronal tumors such as DNETs, tumor resection is the treatment of choice. However, surgery is not always viable due to the difficulties in excising tumors and efficient resection is often limited by unclear margins (Devaux *et al*, 2017). Moreover, despite their benign character, tumor relapse in DNETs and other LGGNTs is not rare, and highly associated to incomplete resection and seizure outbreaks (Chassoux *et al*, 2012; Nolan *et al*, 2004), highlighting a strong need for novel therapeutic approaches. FGFR inhibitors have been tested in clinical trials to treat aggressive cancers harboring alterations in FGFR genes, including cases with FGFR1 alterations, with only partial responses and in some cases severe adverse events (Pant *et al*, 2023; Subbiah *et al*, 2022). Despite this, the spectrum of pan-FGFR and FGFR1-specific inhibitors has increased in the last years, and they have been proposed as a future option for tumors harboring *FGFR1* oncogenic alterations (Fan *et al*, 2024). However, this therapeutic strategy might not be the most efficient in the context of highly stable mutant proteins as our data indicates for N546K and K656E. In line with a minor implication of kinase activity compared to other FGFR1 alterations, a recent study demonstrated that N546K-mutated cells are least sensitive to pan-FGFR inhibitors compared with cell models harboring *FGFR1*-structural variants (Morin *et al*, 2024). Different drug design approaches, targeting protein stability, such as proteolysis-targeting chimeras (PROTACs), an expanding field in oncotherapy (Vikal *et al*, 2024) with already available first candidates targeting specifically FGFR1 (Wang *et al*, 2024), might be taken into consideration for N546K and K656E mutated tumors.

Aside from the two hotspots mutations discussed so far, our work provides extensive data on the susceptibility allele R661P (Rivera *et al*., 2016). FGFR1 R661P-mutant protein clusters close to the WT protein in interactome profiles, suggesting less impact on FGFR1 protein function, compared to the oncogenic mutants. This observation is coherent with the absence of pathological effects other than multinodular DNETs in R661P carriers, who had been clinically followed until adulthood. Most likely in tissues other than brain this variant alone does not lead to major changes in the receptor function, which distinguishes it functionally from other germline *FGFR1* variants that cause developmental conditions. Nevertheless, our results indicate that the R661P variant does have dramatic effects in terms of self-renewal ability and rewiring of gene expression at the target tissue. A similar strong genotype-phenotype specificity at the tissue type-level is observed for germline variants causing osteoglophonic dysplasia and giant cell tumors of the jaw (Gomes *et al*, 2018; White *et al*., 2005). Consistent with the tissue-specificity of the biological effects associated with the R661P variant, differential gene expression analysis revealed enrichment for neural development categories with marked downregulation of key transcription factors involved in glia cell development and oligodendrocyte differentiation, as *SOX9* (Finzsch *et al*., 2008; Hassel *et al*., 2023; Pozniak *et al*, 2010). Notably, a similar association with neuronal transcriptional categories has been recently reported for both *FGFR1*-mutated pediatric low-grade gliomas and neural precursors cell lines (Morin *et al*., 2024).

*FGFR1* is expressed in glial and neural precursors and plays essential functions during brain tissue development and differentiation of neural cell lineages, including oligodendrocyte precursor cells (Furusho *et al*, 2011; Grabiec *et al*, 2016; Yoon *et al*., 2004). In this context, both the oligodendrocyte lineage background and the absence of WT allele expression in R661P HOG cells would contribute to reveal the phenotypic defects caused by the mutant receptor. These changes in FGFR1 activity in neural or oligodendrocyte precursor cells, although compensated by expression of the WT receptor, might prime for the acquisition and selection of activating mutations *in cis* in the oligodendrocytes of these patients. These results may be unveiling a novel model of oncogene-associated susceptibility, based on the need for “correction” of proper signaling and transcriptional programs, which ultimately predisposes toward selection of oncogenic events conferring driving capacity to the mutated clone. This hypothesis is corroborated by the rescue of both stemness/proliferative capacity and expression levels of development-related genes observed in double mutant K656E/R661P and N546K/R661P HOG cells. The R661P mutation in combination with the K656E is also found in one case (likely of sporadic origin based on their variant allele frequency) from the GENIE cohort, arguing that the combination of these two events perpetuates tumorigenesis. Sporadic cases harboring other missense mutations might share common mechanisms with the ones uncovered in the present work, thus enlightening why DNETs and other therapy naïve gliomas with a quiet genome acquire additional hits involving the *FGFR1* gene.

In hereditary tumor syndromes, a continuum model for tumor suppressors evolves the classical two-hit model to a fine-tuning sensitivity of the target cell to either survive, senesce or become tumorigenic depending on the effects of mutational pattern at each tissue level, (Albuquerque *et al*, 2002; Berger *et al*, 2011; Latchford *et al*, 2007). Similarly, a most-optimal level of FGFR1 signaling during early onset tumorigeneis might prevail, with additional layers of complexity due to the key roles of the FGF-FGFR axis plays during development (White *et al*., 2005).

Here we present evidence for modulating effects exerted by *FGFR1* secondary mutations on recurrent hotspot oncogenic mutations, which combined could provide a tolerated level of FGFR1 signaling required during brain development and, on the other hand, trigger oncogenic processes that cause early-onset tumor formation. Future efforts are necessary to confirm this modulatory ability for other missense variants co-identified with the two hotspot mutations. The propensity of certain tumor types to acquire second alterations without any external (treatment) pressure refers to the plasticity of the tumor cells to diverge and adapt through tumor development. Understanding this process might forecast tumor response to external pressures imposed by targeted therapies. The present study provides evidence of novel oncogenic mechanisms and modulatory effects derived from multiple *FGFR1* alterations that need to be considered when choosing treatment options and might help to identify alternative therapeutic approaches for these patients.

## Methods

### Review of public tumor data

Public tumor data was interrogated using the GENIE/cBioportal database (de Bruijn *et al*., 2023) (https://genie.cbioportal.org/), selecting the GENIE Cohort v16.1 (last access in October 2024) and query for *FGFR1* gene. Mutation data was filtered for samples harboring hotspot mutation in one of the codons N546 and K656 and tables with clinical and genomic data was obtained. Data was manually reviewed and second mutations in *FGFR1* were annotated. Mutated samples were divided into brain tumor categories vs others. Mutated samples corresponding to brain tumor types included: Low-Grade Glioma, NOS; Dysembryoplastic neuroepithelial tumor; Pilocytic Astrocytoma; Rosette-forming Glioneuronal Tumor of the Fourth Ventricle; Pilomyxoid Astrocytoma; Miscellaneous Neuroepithelial Tumor; Glioma, NOS; Ganglioglioma; Miscellaneous Brain Tumor, Diffuse Glioma; Glioblastoma Multiforme; High-Grade Glioma, NOS; Anaplastic Oligodendroglioma; Anaplastic Astrocytoma; Diffuse Intrinsic Pontine Glioma; Glioblastoma; Oligodendroglioma; Primary Brain Tumor; Astrocytoma; Encapsulated Glioma). Available data from brain tumor samples was reviewed and tumor grading “low” (for grade 1-2) or “high” (for grade 3-4) was assigned when possible, according to the annotated clinical diagnosis and molecular data following the WHO-IARC Classification of tumors, Central Nervous System, 5^th^ Edition (Louis *et al*., 2021). Tumor cases with tumor grade which was not possible to categorized into low-or high-grade were considered as unknown.

### Cell cultures

The Flp-In T-REx HEK293 (Thermo Fisher ®) cell line was purchased from Thermo Fisher Scientific, while the Human Oligodendroglioma (HOG) cell line was a generous gift from Dr. Lopez Guerrero. Both cell lines have been typified and regularly tested for mycoplasma contamination every two months. Cultures were maintained in Dulbecco’s Modified Eagle Medium (DMEM) with 4.5 g/L glucose (Gibco, Thermo Fisher), supplemented with 10% fetal bovine serum (FBS) (Gibco, Thermo Fisher) and 1% penicillin-streptomycin (P/S) (10 000 U/mL) (Gibco, Thermo Fisher), at 37°C in a 5% CO_2_ atmosphere. Passages were performed once cells reached 80-90% confluency, by trypsinization and subsequent dilution into fresh culture plates.

To induce expression of FGFR1 receptors, Flp-In T-REx HEK293 were treated with 1 µg/mL of Doxycycline (Merck) for 24 hours in every experiment described, except for time course experiment where cells were treated for 8 hours (Fig. 3D-F). For protein stability and degradation experiments, cells were starved overnight with FBS-free DMEM. Next, cells were induced with 25 ng/mL of recombinant human FGF2 (RD Systems) and treated with 100 µg/mL of Cycloheximide (Merck) and 200 nM of Bafilomycin A1 (Merck).

### Generation of stable cell lines and BioID-MS

FGFR1 WT cDNA sequence was recombined into the pDONR-223 vector and then shuttled into the pcDNA5-FRT backbone vector expressing an abortive mutant of BirA (BirA*) tagged with Flag using the gateway recombination cloning system (PMID: 11076863). Stable and inducible Flp-In T-REx HEK293 cell lines expressing FGFR1-BirA*-Flag or Empty-Flag-BirA* or Empty-3xFlag under the tetracycline operator (Tet), were generated according to the manufacturer protocol. Briefly, cells were transfected with the FGFR1-BirA*-FLAG constructs along with a Flippase-expressing plasmid (pOG44), to induce recombination and integration in the cells’ genome. Cells were then selected using 200 µg/mL Hygromycin (Merck) and 5 µg/mL Blasticidin for three weeks. For BioID-MS experiment, cells were seeded in 15 cm plates to reach 70-80 % confluency. The next day, cells were treated with tetracycline (1 μg/µL) and biotin (50 μM, Merck). Cells were then processed for MS analysis as previously described (Hesketh *et al*, 2020). Validation of biotinylation of proximal interactors was carried out through western blot using the anti-strepatavidin-HRP antibody (see western blotting section).

### BioID data analysis

Reproducibility of samples among biological replicates was assessed in R (r-project.org) using the stats package to perform Spearman correlations on spectral counts. To calculate interaction statistics, we used SAINTexpress (Teo et al., 2014) version 3.6.1 on proteins with an iProphet protein probability ≥ 0.99 and unique peptides ≥ 2. Proteomics datasets were compared separately against negative controls (n=22). SAINT analyses^a^ were conducted by compressing n=11 controls (Empty-BirA*-Flag and Empty-3xFlag) to the 11 highest spectral counts among the 22 samples, while baits were not compressed. To infer FGFR1’s high confidence interacting preys, we applied a combination of filters. Preys had to display a SAINT average probability (AvgP) ≥ 0.95, which is more stringent than a BFDR ≤ 0.01 threshold, and an average spectral count (AvgSpec) ≥ 5. Unfiltered contaminants, such as Keratin, BirA*, carboxylases and beta galactosidase were manually removed. We assessed the interaction specificity to each bait by calculating their WD-score with the SMAD R package (Sowa et al., 2009). Heatmaps were created with the R complexheatmap package (Gu et al., 2016). To do so, we first created a matrix containing the AvgSpec of each bait-prey interaction. Unidentified interactions were inputted with an AvgSpec of 0, and a pseudocount of 1 was added to the matrix. The data was log2-transformed, and we calculated Canberra distances between filtered preys and Pearson correlations between baits. Using the Ward. D method, we then extracted 17 clusters of preys and two clusters of baits. Optimal numbers of extracted clusters were estimated on Canberra distances by the silhouette method from the factoextra R package and its fviz_nbclust function. Interaction recalls of FGFR1 were extracted from the human BioGRID interaction database version 4.4.236 (Last access August 2024). Gene ontology (GO) was performed using the clusterProfiler (version 3.0.4) R package, correcting for False Discovery Rate (FDR) through the Benjamini–Hochberg method. GO plots were generated through the ggplot2 (3.5.1) R package.

^a^SAINT scoring was performed on a larger dataset that included localization controls and other mutants not discussed explicitly in the text. All raw data and information on the files included can be found on ProteomeXchange through partner MassIVE MSV000096690.

### Western Blot

Cells were lysed with 100 μL of SDS lysis buffer (25 mM Tris pH 8, 1 mM EDTA, 1% SDS, 10 mM sodium pyrophosphate in dH_2_O). The lysates were incubated at 95°C for 20 min and centrifuged at maximum speed for 10 min. Supernatants were collected and protein concentration was determined using a BCA Protein Assay Kit (ThermoScientific).

For Western blotting, 30 μg of protein was aliquoted and mixed with loading buffer (150 mM Tris pH 6.8, 6% SDS, 15% glycerol and 6% β-mercaptoethanol in dH_2_O). Samples were run on an SDS polyacrylamide gel. Subsequently, the proteins were transferred to a nitrocellulose membrane using the TransBlot Turbo RTA Transfer Kit (Bio-RAD) according to the manufacturer’s protocol. The transferred membranes were blocked in either 5% milk or 5% BSA (depending on the antibody requirements) and incubated with primary antibody overnight at 4°C. The primary antibodies used were anti-FLAG (Merck #F3165), anti-FGFR1 (Abcam #ab76464), anti-phospho-FGFR1 Tyr653/654 (Cell Signaling #52928), anti-PLCγ1 (Cell Signaling #5690), anti-phospho-PLCγ1 Tyr783 (Cell Signaling #2821), anti-tubulin (Merck #F6199) and anti-streptavidin-HRP (Thermofisher #21126). After primary antibody incubation, membranes were washed three times for 5 minutes with TBST buffer (100 mM Tris, 150 mM NaCl and 0.1% tween-20 in dH_2_O) and incubated with HRP-conjugated secondary antibodies at room temperature for 1h. The secondary antibodies used were anti-rabbit-HRP (Merck #A0545) and anti-mouse-HRP (Invitrogen #32430). Then, membranes were washed again three times for 5 minutes with TBST buffer and protein bands were detected using Clarity Western ECL Substrate Kit (BioRad). Images were taken using an Amersham Imager 600 and densitometry analysis was performed using ImageJ software. Three densitometry measurements for each independent experiment have been carried out and outputs have been corrected using loading control (α-Tubulin) values.

### Co-immunoprecipitation

For protein extraction, cells were washed with cold PBS (Corning) and lysed with IP lysis buffer (50 mM HEPES pH 7.4, 150 mM NaCl, 2 mM EDTA, 1% (v/v) NP-40, 30 mM sodium pyrophosphate, 2mM sodium orthovanadate and 0.2% protease inhibitors in dH_2_O) on ice for 30 min. Then, lysates were centrifuged at maximum speed for 30 min and the supernatant was collected. Protein concentration was determined using a BCA assay (ThermoScientific).

Protein G Mag Sepharose beads (Cytiva) were washed in lysis buffer and incubated for 1 hour on rotation with 2 μg/mL of anti-FLAG antibody (Merck #F3165). Following incubation, beads were washed with lysis buffer and incubated with the lysates overnight at 4°C. Then, beads were washed three times with washing buffer (50 mM HEPES pH 7.4, 150 mM NaCl, 2 mM EDTA and 0.1% NP-40), followed by one additional NP-40. Finally, beads were eluted in 30 μL loading buffer and incubated for 10 min at 95°C. The eluted proteins were loaded onto an SDS-PAGE for Western blotting.

### Immunofluorescence staining and imaging

For immunofluorescence staining, cells were fixed in 4% formaldehyde solution for 15 min and permeabilized with 0.05% Triton X-100 for 10 min. Samples were then blocked with 10% goat serum for 1h at room temperature and incubated overnight at 4°C with the primary antibodies anti-FGFR1 (Abcam #ab76464) and anti-α-tubulin (Merck #F6199). After primary antibody incubation, samples were washed and incubated for 1h at 37°C with the fluorophore-conjugated secondary antibodies anti-mouse Alexa Fluor 488 (Invitrogen #A21202) and anti-rabbit Alexa Fluor 555 (Invitrogen #A21430). Finally, samples were mounted with Prolong Gold Antifade Reagent with DAPI (Invitrogen), and images were captured using a Leica SP5 inverted confocal microscope.

### CRISPR/Cas9 genome editing

The coding sequences for single-guide RNAs designed to target specific *FGFR1* loci have been cloned into the PX458 vector (Addgene #48138) as described in (Ran *et al*, 2013) and delivered through lipofection in HOG cells together with specific synthetic donor single strand DNAs (Alt-R™ HDR Donor Oligos, Integrated DNA Technologies). Donor templates were designed to introduce additional silent point mutations for PAM silencing and restriction enzyme-based screening (Fig. EV6A). Cells were sorted in multiwell96 plates as single cells per well and colonies were identified after two weeks. To evaluate sgRNAs efficacy, test based on cleavage with T7 endonuclease (New England Biolabs) have been carried out prior to experiment. Clones have been grown and screened through PCR followed by cleavage with specific restriction enzymes and editing has been confirmed by Sanger sequencing.

### Clonogenic assays

For 2D clonogenic assays, cells were seeded in 6-well plates at a density of 10^3^ cells per well in fresh medium and incubated for 2 weeks. After incubation, colonies were fixed with methanol for 10 min and stained with 0.1% (w/v) crystal violet in dH_2_O for 20 min. Plates were then washed with dH_2_O to remove the excess of stain crystal and allowed to dry. Stained plates were scanned, and colonies were quantified using ImageJ software.

For 3D colony formation experiments in Matrigel, 100 μL of Matrigel (Corning) was added to 1 cm^2^ wells and led solidify for 15 min at 37°. Subsequently, 1×10^4^ cells were seeded into each well and incubated for 1 week at 37°C in a 5% CO_2_ atmosphere. Images were captured using a Carl Zeiss™ Axio Vert.A1 microscope (Zeiss Zen software) and colony area was quantified for colonies with a minimum size of 100 μm^2^ using ImageJ software. Each condition was plated in duplicate (2 wells) and three independent fields were analyzed for each well.

### Quantitative RT-PCR

RNA was extracted from the cells using the RNeasy kit (Qiagen), and RNA concentration was determined using a Nanodrop One spectrophotometer. cDNA was synthetized using the SuperScript Reverse Transcriptase II kit (Invitrogen) and quantitative PCR reactions were performed using the PowerUp SYBR Green Master Mix (Applied Biosystems) on a LightCycler 480 thermocycler. The data was analyzed using the 2^-ΔΔCt^ method, calculating mean ± SEM using three replicates per sample and correcting for housekeeping genes (*HPRT1* for HEK293 cells and *ACTB* for HOG cells) expression values.

### 3D protein structure prediction

In the absence of a crystal structure of the full FGFR1-PLCγ protein complex in the Protein Data Bank, we employed AlphaFold3 (AF3, developed by DeepMind (Abramson *et al*., 2024)) to generate an approximate model of the complex. This predictive approach allowed us to explore the potential location of the binding interaction between PLCγ and the FGFR1 kinase domain, yielding high-confidence metrics for both the interfacial predicted modeling (ipTM = 0.45) and the predicted template modeling (pTM = 0.54), with pIDDT values above 70 for residues located at the interface.

The selected models were refined using the QuickPrep protocol in MOE2020 (Molecular Operating Environment (MOE), 2024). The AlphaFold3 model for FGFR1 (*Homo sapiens*, UniProt ID: P11362) shares 99.7% similarity with the crystallized human active form (PDB ID: 3GQI), achieving 77.3% overlap. Structural alignment of the model with the original pocket residues displayed a strong superposition, with an average RMSD of 4.81 Å (Fig. EV5). Overall, the resulting complex FGFR1-PLCγ predicted by AlphaFold3 agrees with previously reported experimental data by Hajicek and colleagues (Hajicek *et al*., 2019).

### RNA library construction and sequencing

RNA samples were collected in triplicates using the RNeasy kit (Qiagen) and integrity was evaluated using RNA 6000 Nano Assay on a Bioanalyzer 2100 (Agilent). Libraries were generated using the TruSeq® Stranded mRNA LT Sample Prep Kit (Illumina Inc., Rev.E, October 2013). Libraries were sequenced by the standard Illumina protocol to create raw sequence files (.fastq files), which underwent quality control analysis using FastQC. To avoid low-quality data negatively influencing downstream analysis, we trimmed the readson the 3′-end and only used the first 51 bp from the 5′-end of each read for further analysis.

### Analysis of RNA-seq data

RNA-seq reads were mapped against the human reference genome (GRCh38) with STAR 2.7.8a (Dobin *et al*, 2013) using ENCODE parameters. Genes were quantified with RSEM 1.3.0 (Li & Dewey, 2011) using the Gencode v44 annotation. Genes with at least 1 count-per-million reads (cpm) in at least 3 samples were kept. Differential expression analysis was performed with limma R package v3.54.2 (Ritchie *et al*, 2015) using TMM normalization. The voom function (Law *et al*, 2014) was used to transform the count data into log2-counts per million (logCPM), estimate mean-variance relationship and to compute observation-level weights. These voom-transformed counts were used to fit the linear models. Volcano plots were generated using the ggplot R package. Gene ontology (GO) was performed using the clusterProfiler (version 3.0.4) R package, correcting for False Discovery Rate (FDR) through the Benjamini–Hochberg method. Cutoffs for significant differentially expressed genes were set at Log2FC <-0.7 or > 0.7 and FDR ≤ 0.05. GO plots were generated through the ggplot2 (3.5.1) R package.

### Statistical analysis

Correlation analysis between groups has been performed through a two-sided Chi-square test with a confidence interval of 95%. Data obtained from cellular assays and molecular analysis was tested using Analysis of Variance (ANOVA) or Krustal-Wallis method (when normality was not assumed). For multiple comparisons, a post hoc pairwise comparisons analysis was conducted using the Tukey Honest Significant Difference (Tukey HSD) test for ANOVA or Dunn’s test for Krustal-Wallis test, to assess the significant differences between groups. All statistical tests were performed at an adjusted significance level of p <0.05. The statistical analysis was performed using R (R Core team 2022, R version 4.3.3).

## Supporting information

BioID-MS_Datasets

RNAseq_Datasets

## Acknowledgements

We would like to thank Prof William Foulkes for the support along all stages of this work. Prof. Lopes-Guerrero and Dr Antonio Gentinella for the kind gift of the HOG cell line and the Streptavidin-HRP conjugate reagent, respectively. Dr Meritxell Rovira and Dr Cristina Muñoz for sharing antibodies and reagents. We would like to thank also Dr Tenzin Gayden for the help in the early stages of this conceptual work and the MNFC peer mentoring group. Dr Sidong Huang, Dr Ruth Rodriguez and Dr Marta Pineda for helpful discussion. Finally, we also acknowledge Helena Martí for helping with figure making.

## Author contributions

**Jacopo Boni:** Conceptualization, Data curation, Formal analysis, Investigation, Methodology, Visualization, Writing – original draft, Writing – review and editing. **Míriam Fernández-González:** Data curation, Formal analysis, Investigation, Methodology, Visualization, Writing – original draft. **Hyerim Han:** Investigation, Methodology, Writing – review and editing. **Carla Roca:** Data curation, Formal analysis, Writing – review and editing. **Cassandra J. Wong:** Investigation, Methodology. **Cristina Rioja:** Formal analysis, Writing – review and editing. **Clara Nogué:** Investigation, Methodology, Visualization. **Leticia Manen-Freixa:** Formal analysis. **Jonathan Boulais:** Formal analysis, Software.

**Endika Torres-Utizberea:** Formal analysis. **Antonio Gomez:** Data curation, Software. **Martin Haselblatt:** Investigation, Writing – review and editing. **Roger Estrada-Tejedor:** Supervision. **Albert A. Antolin:** Software, Supervision. **Islam E. Elkholi:** Data curation, Writing – review and editing. **Nada Jabado:** Supervision. **Jean-François Côté:** Supervision. **Anne-Claude Gingras:** Supervision. **Barbara Rivera:** Conceptualization, Investigation, Funding acquisition, Project administration, Project supervision, Writing – review and editing.

## Expanded view Figures

**Expanded view Figure 1.**
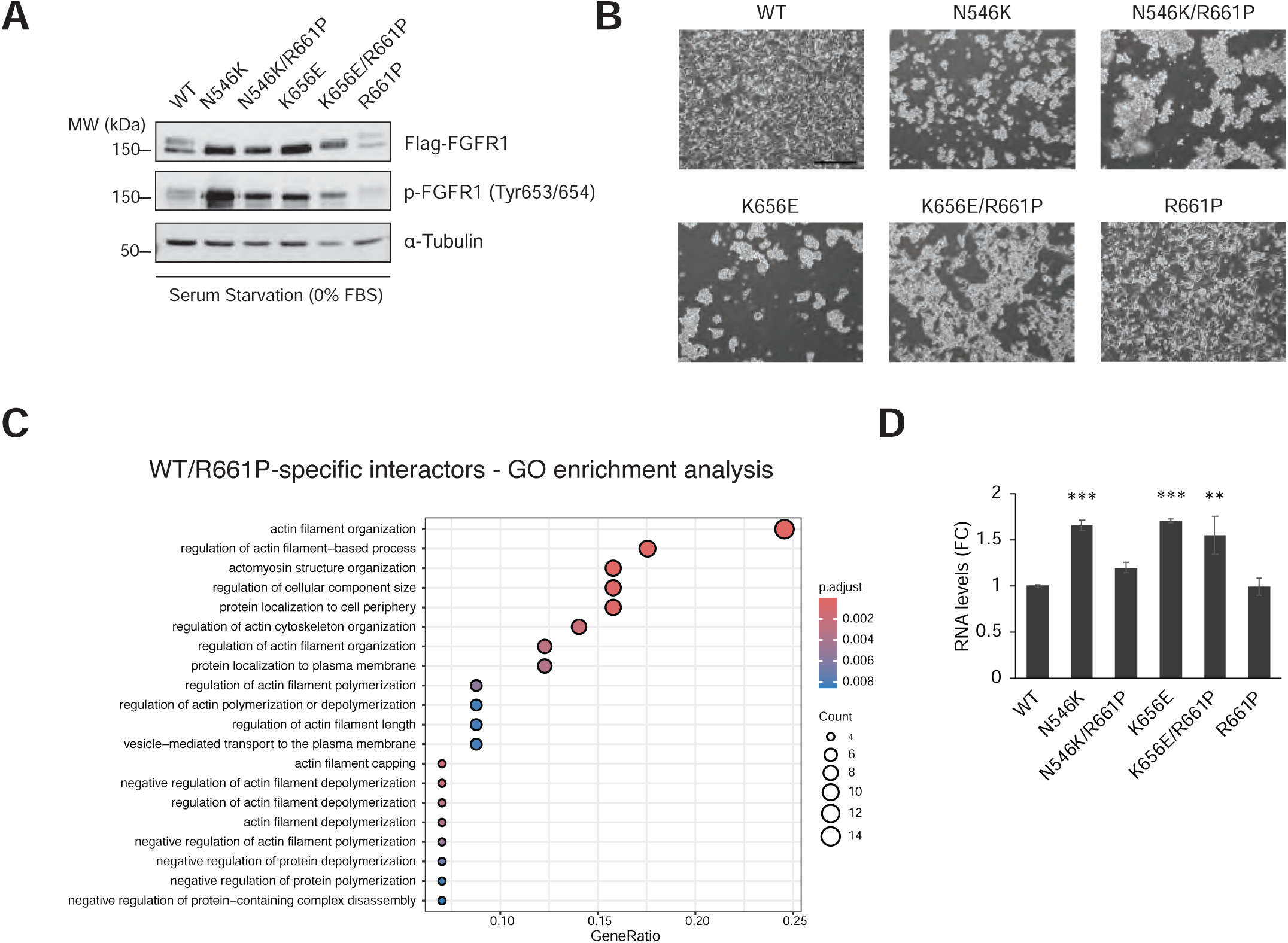
A) Western blot of the six cell lines used in the present study, showing expression and activation (phosphorylation of Tyr 653/654) of FGFR1, after 24h of Tet induction and serum starvation (0% FBS overnight). **B)** Optical microscope captures of T-REx Flp-In HEK293 expressing WT and mutant FGFR1 proteins 24 hours post Tet-induction. Scale bar: 200 µm. **B)** Dot plot of GO enrichment analysis using the list of proteins found in Clusters 14-15 of the heatmap in Fig. 2H (R661P-specific + WT/R661P shared interactors). Size and color of dots represent numbers of matched genes and adjusted p-value (p.adjust), respectively. **D)** FGFR1 RNA levels in WT and the five mutant T-REx Flp-In HEK293 cell lines, obtained by RT-qPCR. Data is represented by Mean ± SEM from n =2 independent experiments and significant differences against WT condition have been indicated (ANOVA test, p value * ≤ 0.05, ** ≤ 0.01, *** ≤ 0.001). GO, Gene Ontology.

**Expanded view Figure 2.**
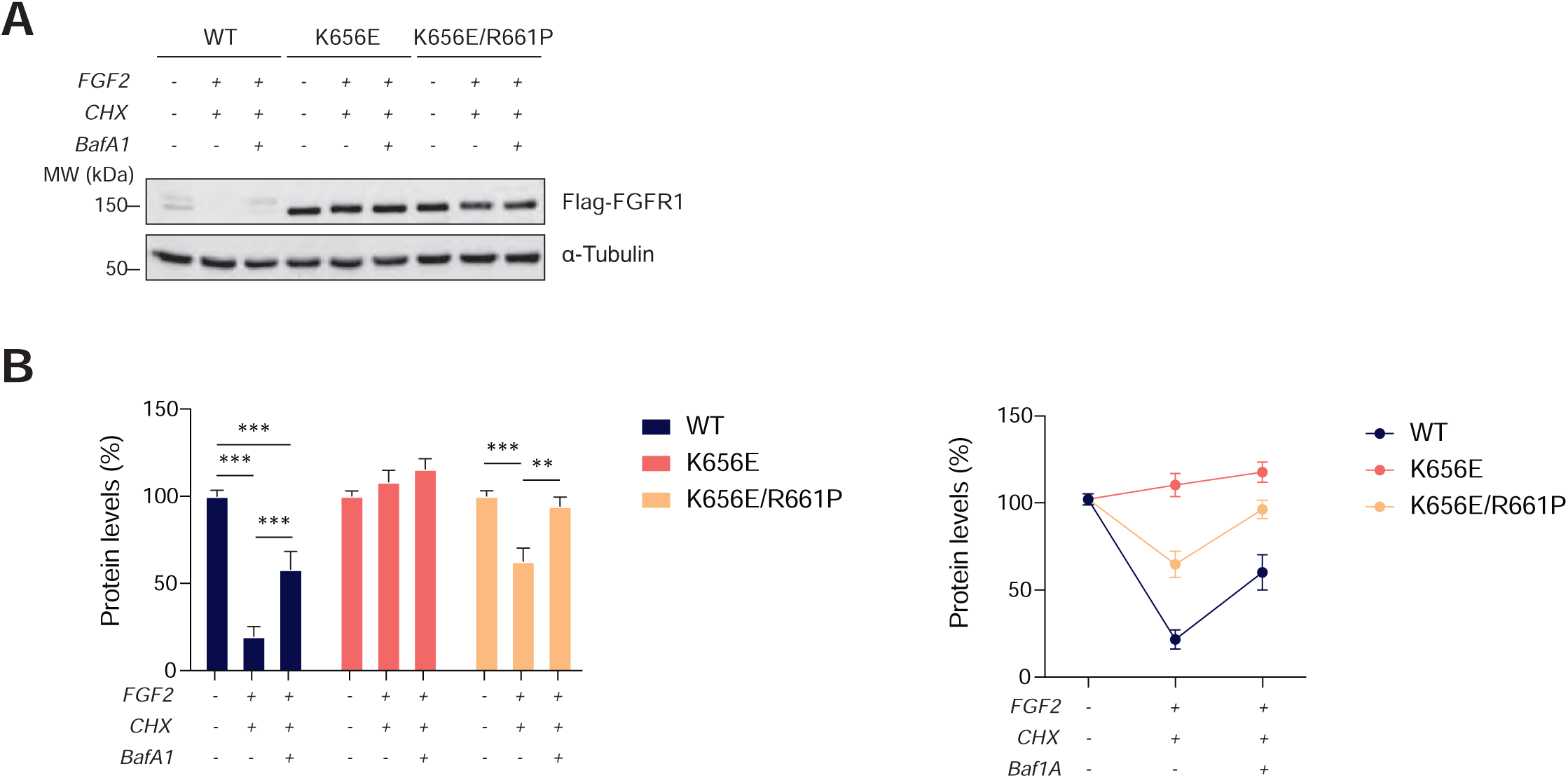
A) Western blot analysis of WT FGFR1, K656E and K656E/R661P mutants indicating total Flag-FGFR1 protein levels upon FGF2 induction and treatment with only cycloheximide (100 µg/mL) or cycloheximide combined with Bafilomycin A1 (200 nM) for 6h. **B)** Relative amounts of WT, single and double mutant FGFR1 protein obtained by quantifying western blots from n = 4 independent experiments. Values have been normalized (FC) against each relative 0h reference values. Data is represented by mean ± SEM and significant variations in protein amounts against each specific reference values have been indicated (ANOVA test, p value * ≤ 0.05, ** ≤ 0.01, *** ≤ 0.001) in the bar plots.

**Expanded view Figure 3.**
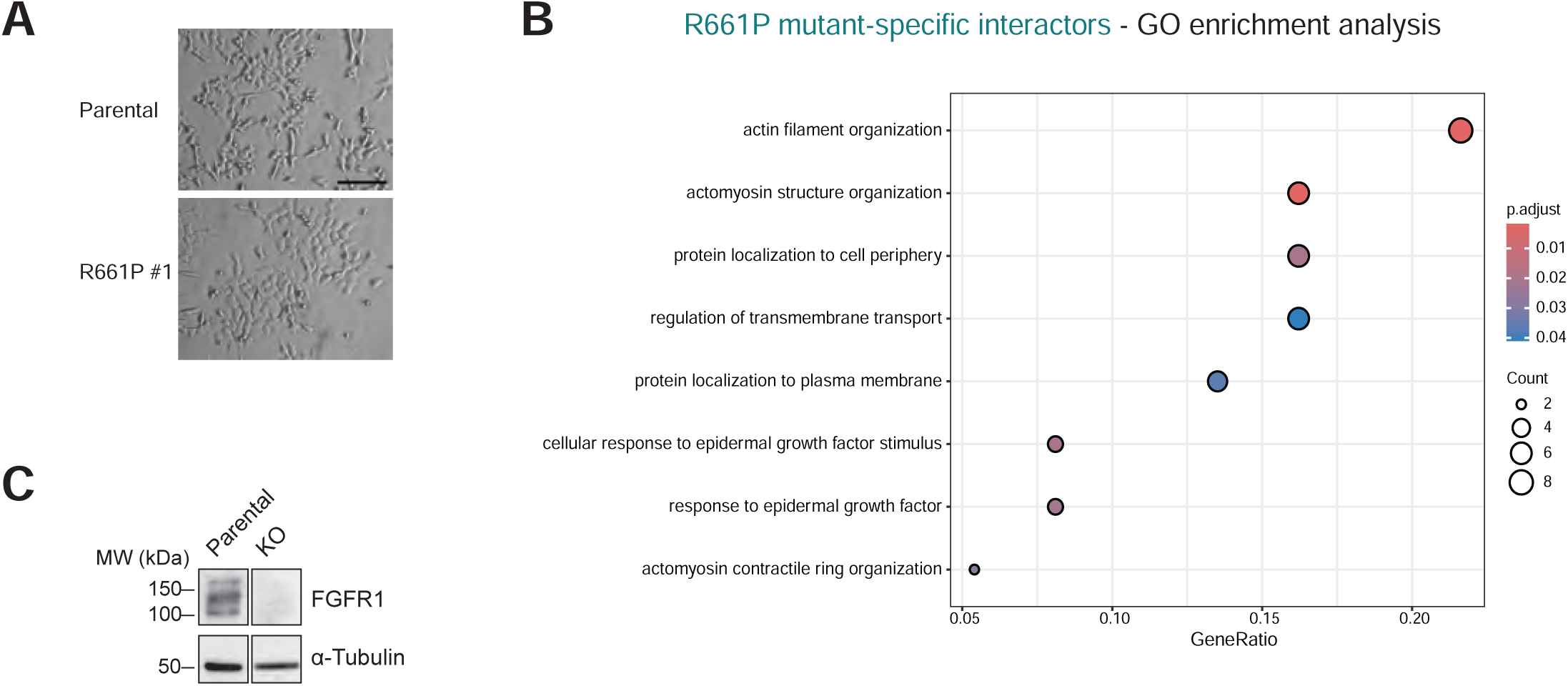
A) Optical microscope captures of Parental (upper panel) and R661P #1 (lower panel). Scale bar: 200 µm. **B)** Dot plot of GO enrichment analysis using the list of preys forming Cluster 14 of the heatmap in Fig. 2H (BioID R661P-specific interactors). Size and color of dots represent numbers of matched genes and adjusted p-value (p.adjust), respectively. **C)** Western blot of FGFR1 protein confirming absence of FGFR1 expression in KO HOG cells, compared to Parental control. GO, Gene Ontology.

