## Supplementary material and figures for "Multiple *FGFR1* mutations modulate tumorigenic mechanisms in glioneuronal tumors"

Title: **Modelling multiple *FGFR1* mutations uncovers modulatory effects and novel tumorigenic mechanisms in glioneuronal tumors.**

**Appendix – Table of Contents**

**Supplementary Figures and Tables**

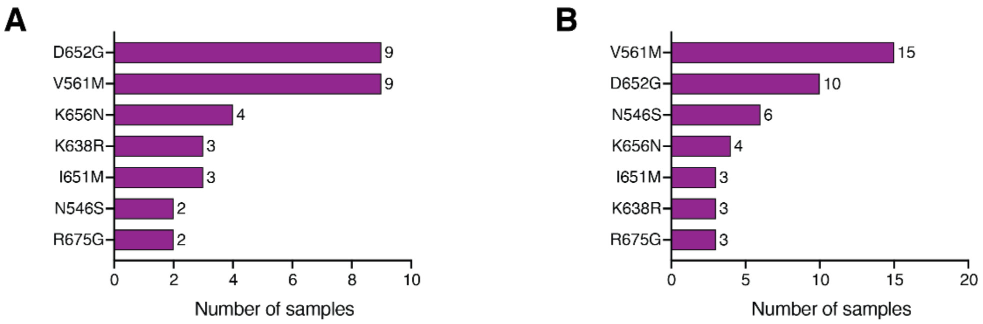

**Appendix Figure S1.** Most recurrent secondary missense variants in FGFR1 genes identified in tumor samples from the GENIE database, co-occurring with one of the hotspots N546K/K656E. Number of cases for each mutation (only mutations appearing in at least n = 2 cases have been plotted).

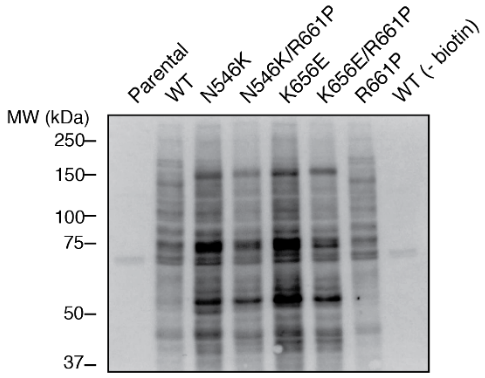

**Appendix Figure S2.** Western blot showing the profiles of biotinylated proteins for the six baits, 24 hours post Tet-induction and biotin treatment, detected through streptavidin-HRP. Parental cell line and WT FGFR1-BirA*-Flag with no biotin added have been used as controls.

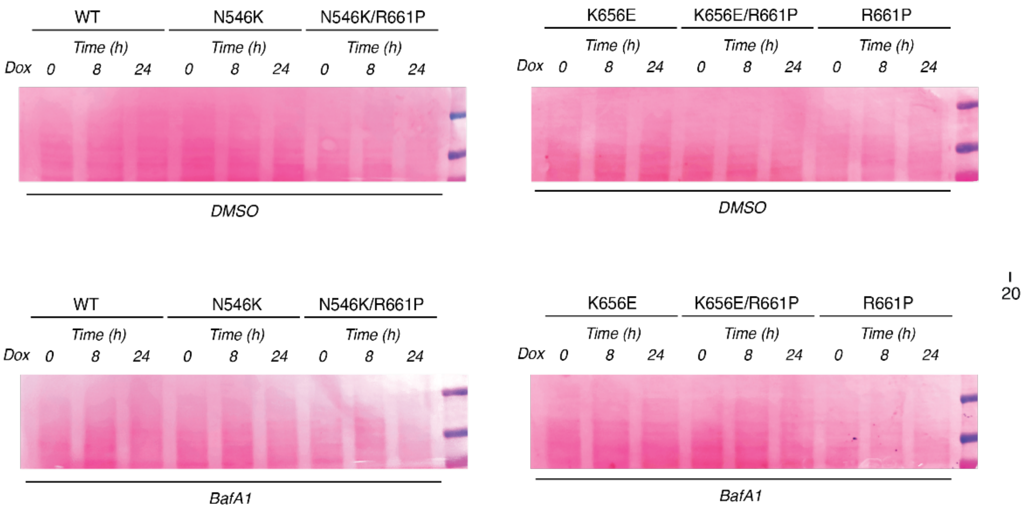

**Appendix Figure S3.** Ponceau staining of western blot membrane of the time course experiment illustrated in Fig. 3F.

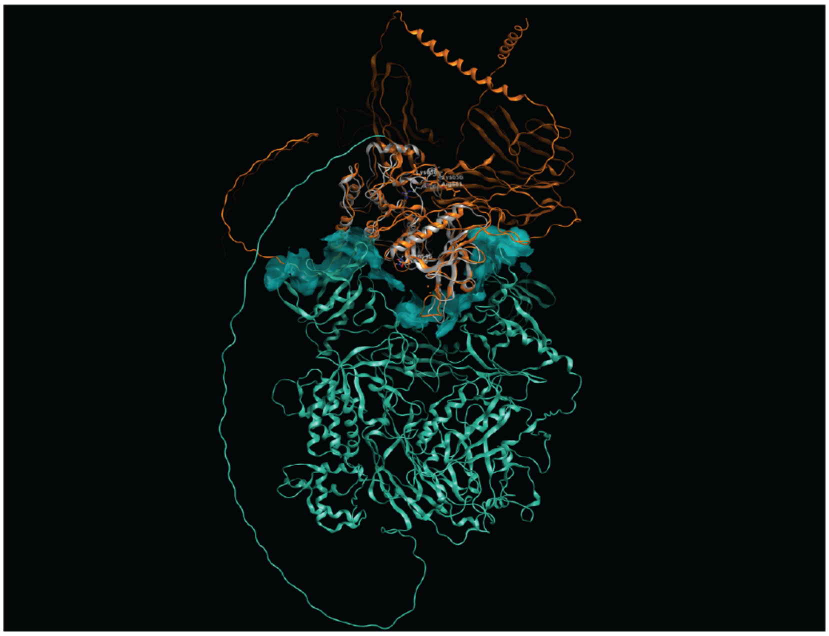

**Appendix Figure S4.** Overlap of the crystallized human active form of the kinase domain of FGFR1 protein 3GQI and AlfaFold 3 FGFR1-PLCγ complex. In orange; predicted FGFR1 protein by AF3. In light grey: crystallised active FGFR1 kinase domain 3GQI. In light blue: predicted PLCγ protein by AF3.

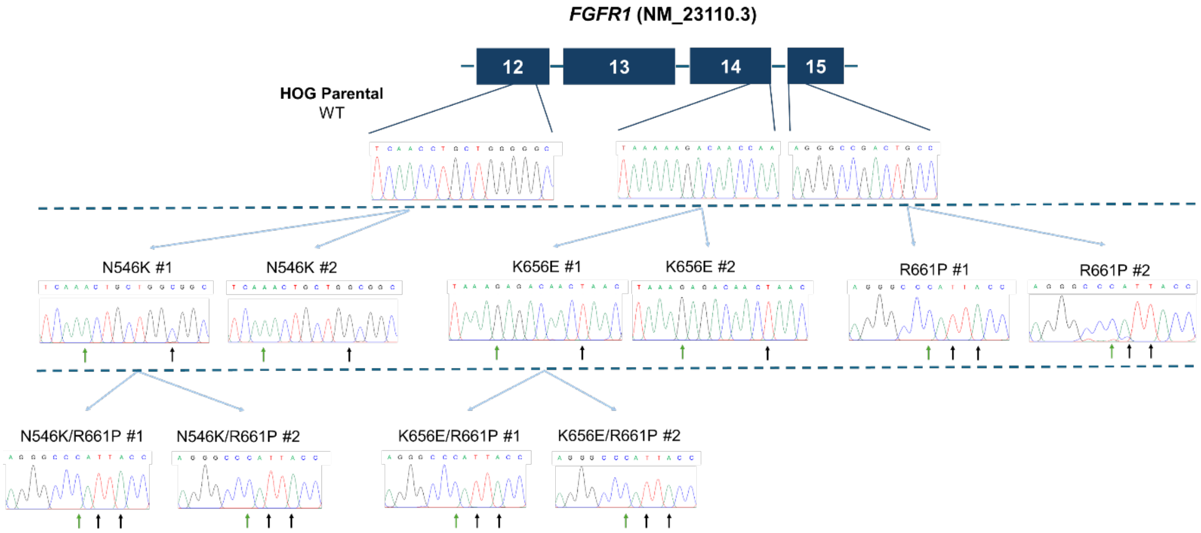

**Appendix Figure S5.** CRISPR/Cas9-editing of HOG cell lines was performed in multiple steps. Single FGFR1 mutant N546K, K656E and R661P were generated in a first step. R661P editing was performed to generate double mutant clones using N546K #1 and K656E #1 clones as parental cell lines. Correct genome editing was assessed through PCR and Sanger sequencing. Blue arrows indicate SNV causing the desired missense mutations, while black arrows indicate additional silent mutations (PAM and/or restriction sites).

| N546K | |
| --- | --- |
| **Amino acid change** | **Number of patients** |
| N546K | 213 |
| N546D | 24 |
| N546S | 6 |
| N546H | 1 |
| Total | 244 |

| K656E | |
| --- | --- |
| **Amino acid change** | **Number of patients** |
| K656E | 121 |
| K656N | 11 |
| K656M | 4 |
| K656Q | 1 |
| K656D* | 3 |
| Total* | 137 |

**Appendix Table S1.** Distribution of each missense variants identified at FGFR1 codon N546 (N546K, N546D, N546S and N546H) and K656 (K656E, K656N, K656M, K656Q and K656D) in patients from the GENIE cohort. *K656D is most likely a combination of the missense variants K656N and K656E and therefore K656D cases have been counted also in the K656E and K656N categories.

### Supplementary Methods

| Primer | Forward | Reverse |
|---|---|---|
| ***FGFR1* Exon 14-15** | CGCTTGCTGTGATGAGAAGCCTG | GCCTTTCAACATCTGGAGCAGAG |
| ***FGFR1* Exon 12** | CCCACTCCCTTAGCCTTTATCC | CTCTTAACCCCCTTCCCTAGC |
| ***FGFR1* Exon 4-5** | TGTCCGTGTTCATCTGGAACTG | TGAAAAGCATGTAATCAGGACTTC |

**Appendix Table S2.** Primer sequences for PCR.

| Primer | Forward | Reverse |
|---|---|---|
| ***SOX9*** | GGCAAGCTCTGGAGACTTCTG | CCCGTTCTTCACCGACTTCC |
| ***GLI2*** | TTTGTCTCTCTCGGATTGCCA | AGGAGAGGCCTTTTTACCCG |
| ***HIF1A*** | AAGCCTTGGATGGTTTTGTTATGG | CCCTTTTTCACAAGGCCATTTCT |
| ***WNT5A*** | GCTTTGCCAAGGAGTTCGTG | CCAGGTTGTACACCGTCCTG |
| ***NRP1*** | TGTGAAGTGGAAGCCCCTAC | TGGTGCTGTCTATGACCGTG |
| ***SEMA3E*** | CCGGTTACGCCTGTCACATA | ACTCGGCCAGTGTATCTCTT |
| ***THBS1*** | CAGGAGCAACCTCTACTCCG | CAGCAGGGATCCTGTGTGT |
| ***FN1*** | CACCTGTACCCACACGGTC | TCCAGGAACCCTGAACTGTAAG |

**Appendix Table S3.** Primer sequences for qPCR.

| CRISPR | Forward | Reverse |
|---|---|---|
| **N546K** | CACCGAAGAATATCATCAACCTGCT | AAACAGCAGGTTGATGATATTCTTC |
| **K656E** | CACCGGCCTTGTCGGCACTCACGT | AAACACGTGAGTGCCGACAAGGCC |
| **R661P** | CACCGCCATCCACTTCACAGGCAGT | AAACACTGCCTGTGAAGTGGATGGC |
| **KO** | CACCGAGATGCTCTCCCCTCCTCGG | AAACCCGAGGAGGGGAGAGCATCTC |

**Appendix Table S4.** Guide RNA sequences targeting FGFR1.

| CRISPR | Sequence |
|---|---|
| **N546K** | AGATGATGAAGATGATCGGGAAGCATAAGAATATCATCAAACT GCTGGCGGCCTGCACGCAGGATGGTGGGTGCCGGCCAGAC |
| **K656E** | AGATGAAACCACCAGCACAGGGCGGCCTTGTCGGCACTCACGTT AGTTGTCTC TTATAGTAGTCGATGTGGTGAATGTCCCGTGCGAGGCCA |
| **R661P** | CCGGTCAAATAATGCCTCGGGTGCCATCCACTTCACAGGTAAT GGGCCCTGAAAGCAGCACAGGGGAGGTTGGAGTGGCCCCAG |

**Appendix Table S5.** Donor template sequences.
